# Rostro-caudal specificity of corticospinal tract projections in mice

**DOI:** 10.1101/2020.07.21.214908

**Authors:** Oswald Steward, Kelly M. Yee, Mariajose Metcalfe, Rafer Willenberg, Juan Luo, Ricardo Azevedo, Jacob H. Martin-Thompson, Sunil P. Gandhi

**Author notes:** Current address: Department of Neurology, Cedars-Sinai Medical Center, Los Angeles, CA 90048. Corresponding Author: Oswald Steward, Ph.D., 1101 GNRF, 837 Health Sciences Dr. University of California at Irvine, Irvine, CA 92697.

## Abstract

Rostro-caudal specificity of corticospinal tract (CST) projections from different areas of the cortex was assessed by retrograde labeling with fluorogold and retrograde transfection following retro-AAV/Cre injection into the spinal cord of tdT-reporter mice. Injections at C5 led to retrograde labeling of neurons throughout forelimb area of the sensorimotor cortex, the rostral forebrain area (RFA), and a region in the lateral cortex near the barrel field. Injections at L2 led to retrograde labeling of neurons in the posterior sensorimotor cortex (hindlimb area) but not the RFA or lateral cortex. With BDA injections into the main sensorimotor cortex (forelimb region), labeled axons terminated selectively at cervical levels. With BDA injections into caudal sensorimotor cortex (hindlimb region), labeled axons passed through cervical levels without sending collaterals into the gray matter and then elaborated terminal arbors at thoracic-sacral levels. With BDA injections into the RFA and lateral cortex near the barrel field, labeled axons terminated at high cervical levels. Axons from medial sensorimotor cortex terminated primarily in intermediate laminae; Axons from lateral sensorimotor cortex terminated primarily in deep layers of the dorsal horn. One of the descending pathways seen in rats (the ventral CST) was not observed in most mice.

**SIGNIFICANCE:** Mice are used extensively for studies of regeneration following spinal cord injury because of the ability to create genetic modifications to explore ways to enhance repair and enable axon regeneration. A particular focus has been the corticospinal tract (CST) because of its importance for voluntary motor function. Here, we document features of the rostro-caudal specificity of CST

## INTRODUCTION

Mice are used extensively for studies of regeneration following spinal cord injury because of the ability to create genetic modifications to explore the role of particular genes in regulating or enabling axon regeneration (Zheng et al 2006). Regeneration of the corticospinal tract (CST) has been of particular interest because of the key role the CST plays in controlling voluntary motor function. Recent studies have documented that several different genetic manipulations can enable regeneration of CST axons past a spinal cord injury (Du et al 2015, Hollis et al 2016, Liu et al 2010, Wang et al 2015) and/or promote sprouting of uninjured CST axons after partial injury (Geoffroy et al 2015).

One question that has not been addressed in most studies is the degree to which regenerated CST axons recapitulate normal specificities of projection. For example, do the CST axons that extend past a cervical level injury originate from neurons that normally project to cervical levels or are they axons that normally project to lumbar and sacral levels? The answer is important for interpreting recovery of motor function and cortical remapping that occurs in conjunction with regenerative growth (see for example Hollis et al., 2016).

Addressing questions about rostro-caudal specificity of regenerated CST axons and interpreting accompanying recovery of motor function requires an understanding normal CST projection specificity. In this regard, some aspects of projection specificity to spinal levels (C7 vs. L4) have been explored (Kamiyama et al 2015), but details of projection specificity from different parts of the sensorimotor cortex have not been extensively explored in mice.

The primary goal of the present study was to map the rostro-caudal projection patterns from different parts of the sensorimotor cortex to the spinal cord in mice. We were especially interested in determining whether CST axons that project from hindlimb cortex to lumbo-sacral sections also gave rise to collaterals at cervical levels and whether CST axons from forelimb regions also extended axons to lumbar levels. Another aspect of CST organization that is not well-understood is the pattern of projection of neurons outside the main sensorimotor cortex. Injections of retrograde tracers into the cervical spinal cord of rats leads to retrograde labeling of pyramidal neurons in the rostral forelimb area (RFA) and in an area in the dorso-lateral neocortex cortex near the barrel field (Nielson et al., 2010) but projection patterns of neurons in these regions is unknown. The other aspect of CST organization that is not well-studied is the degree of bilaterality of CST projections from different parts of the sensorimotor cortex. This is of interest because trans-midline sprouting after unilateral pyramidotomy is one of the main models for testing interventions to enhance sprouting after injury (Geoffroy et al 2015).

Here, we assess topography of CST projections by mapping the cells of origin of CST projections and by mapping projections from different areas using BDA tract tracing. Key findings are: CST axons from the forelimb region of the main sensorimotor cortex terminate selectively in cervical levels without extending to thoracic levels and below; 2) Axons from the caudal sensorimotor cortex (hindlimb region) extend through cervical regions to terminate in lumbar through sacral segments without extending collateral branches into the cervical gray matter; 3) The rostral forelimb area in the frontal cortex and the lateral neocortex near the barrel field project selectively to high cervical levels; 4) Axons from the medial portion of the main sensorimotor cortex (M1) arborize primarily in the intermediate lamina with some axons extending into the ventral horn whereas axons from the lateral portion of the main sensorimotor cortex arborize primarily in the dorsal horn; 5) The degree of bilaterality of CST projections varies by area and level.

## METHODS

### Experimental animals

These studies involved mice of different strains (Table I): Balb/C, were purchased from Harlan Labs, Rosa^tdTomato^ mice with a lox-P flanked STOP cassette that prevents expression tdTomato (obtained originally from Jackson Labs and maintained in our breeding colony for several generations), *Pten*^*loxP/loxP*^ /Rosa^tdTomato^ transgenic mice that we generated by crossing *Pten*^*loxP/loxP*^ with Rosa^tdTomato^ mice (maintained in our breeding colony) and CST-YFP mice (Bareyre et al 2005), which are the F1 progeny of a parental mouse with *Emx-Cre* and a mouse with *Thy1-stop-YFP* [B6.Cg-Tg(Thy1-EYFP)15Jrs/J, Jackson Labs]. Rosa^tdTomato^ mice and CST-YFP mice have a C57Bl/6 genetic background; *Pten*^*loxP/loxP*^ /Rosa^tdTomato^ transgenic mice are mixed genetic background (C57Bl/6 & 129Sv).

**Table I:** Strain, tracer type, target coordinates, and animal number The table lists the mouse strain for each experiment, AAV vector type, type of BDA used (mini-ruby conjugated or un-conjugated), target coordinates for the injections and animal number. All cortical injection depths are 0.5mm unless otherwise noted. Both Rosa^tdTomato^ and PTEN^f/f^;Rosa^tdTomato^ mice are included because the accompanying PTEN deletion in PTEN^f/f^;Rosa^tdTomato^ mice is considered to be irrelevant for this study.

| Strain and number | Tracer | Target coordinates | Animal number |
| --- | --- | --- | --- |
| $Rosa^{tdTomato}$<br>$PTEN^{ff};Rosa^{tdTomato}$<br>(n=8) | retro-AAV/Cre | C5 (n=4)<br>L2 (n=4) | 0315A-H<br>1month survival |
| $Rosa^{tdTomato}$<br>N=2 | retro-AAV/Cre | L2 | 0612A,B<br>3 week survival |
| $PTEN^{ff};Rosa^{tdTomato}$<br>(n=2) | retro-AAV/Cre<br>at L2, retro-AAV/shPTEN-GFP at C5 | C5 and L2 | 0612C,D<br>3 week survival |
| $Rosa^{tdTomato}$<br>$PTEN^{ff};Rosa^{tdTomato}$<br>(n=11) | retro-AAV/Cre | C5 (n=8)<br>L2 (n=3) | 0820A,B,L,M,N,O,P<br>1 year survival |
| CST-YFP, n=3 | Fluorogold | C5 spinal<br>cord 0.5L |  |
| Balb/C, n=3 | BDA<br>unconjugated | 1.0A, 1.5L | 1120A,B,C |
| Balb/C, n=5 | BDA miniruby | 0.5A, 1.0L | 0225E,F,G,H,I |
| Balb/C, n=3 | BDA miniruby | 1.0A, 1.0L | 0225A,B,D<br>0321A,B,C |
| Balb/C, n=3 | BDA<br>unconjugated | 0-at bregma,<br>1.0L | 1120D,E,F |
| Balb/C, n=3 | BDA<br>unconjugated | 1.0P, 1.0L | 1120G,H,I |
| Balb/C, n=6 | BDA miniruby | 1.0P, 1.5L | 0225J-O |
| Balb/C, n=6 | BDA miniruby | 2.0P, 1.5L | 0225P<br>0321E-I |
| Balb/C, n=3 | BDA miniruby | 2.8A, 0.8L | 0313A-C |
| Balb/C, n=3 | BDA miniruby | 1.1P, 3.2L,<br>1.1 deep | 0408A-C |
| Balb/C, n=3 | BDA<br>unconjugated | 1.0A, 2.25L | 0812A-C |
| Balb/C, n=3 | BDA<br>unconjugated | 0-at bregma,<br>2.25L | 0812D-F |
| Balb/C, n=3 | BDA<br>unconjugated | 1.0P, 2.25L | 0812G-I |

### Fluoro-Gold Injections for retrograde labeling

To generate a map of cortical neurons that give rise to the CST, three adult CST-YFP mice were injected bilaterally with a total of 0.4 μl of 4% Fluoro-Gold (Fluorochrome, LLC; Denver CO) at cervical level 5 (C5). Mice were anesthetized with isofluorane, and the spinal cord was exposed by laminectomy at C5. Injections were made bilaterally at 0.5mm lateral to the midline and 0.5mm deep, with 0.2μl injected at each site using an electronically-controlled injection system (Nanoliter 2000 injector and Micro4 pump controller, World Precision Instruments) fitted with a pulled glass micropipette. One week following injections mice were transcardially perfused with 4% paraformaldehyde in 0.1M phosphate buffer (4% PFA), and brains and spinal cords were dissected and post-fixed in 4% PFA overnight before being equilibrated and stored in 27% sucrose at 4°C.

Brains from mice injected with Fluoro-Gold were embedded by freezing in TissueTek O.C.T. (VWR International). Cryostat sections were taken in the coronal plane at 30μm, and free-floating sections were collected in PBS at 420μm intervals through the cortex and mounted onto gelatin-coated slides. Brain sections were imaged at 4X on an Olympus IX80 fluorescence microscope, and overlapping images were stitched together in ImageJ (Rasband, W.S., ImageJ, National Institutes of Health, http://imagej.nih.gov/ij/, 1997-2014) using linear blending fusion in the Preibisch stitching plugin (Preibisch et al 2009).

### Retrograde transfection of cortical motoneurons with retro-AAV/Cre in tdTomato reporter mice

To label cortical motoneurons in a way that allows visualization of dendritic arbors and produces labeling that is optimal for light sheet microscopy, we used AAVs that are transported retrogradely transported (retro-AAV) in transgenic Rosa^tdTomato^ and *Pten*^*loxP/loxP*^ /Rosa^tdTomato^ mice. Adult mice received bilateral injections of retro-AAV/Cre (0.3μl each) at cervical level 5 (C5) or L2 vertebral level (see Table I for a summary of cases). Mice were anesthetized with isofluorane, and the spinal cord was exposed by laminectomy at vertebral level C5 or L2. Injections were made using a Hamilton Three-four weeks post-injection (see Table I), mice were transcardially perfused with 4% paraformaldehyde in 0.1M phosphate buffer (4% PFA), and brains and spinal cords were dissected, post-fixed in 4% PFA overnight, and stored in buffer at 4°C. Another set of mice received injections at C5 (n=8) or L2 (n=3) and survived for 1 year post-AAV injection to assess whether there were any negative consequences of long-term PTEN deletion via retro-AAV/Cre.

To assess whether there were populations of CMNs that could be co-transfected by retro-AAV injected into different spinal levels (C5 vs. L2), mice (n=2) received injections of retro-AAV/Cre into the lumbar spinal cord (L2 vertebral level) and retro-AAV/shPTEN-GFP (a vector we engineered to knock down PTEN for studies of axon regeneration in rats).

### Fixation and tissue processing

Mice were perfused trans-cardially with 4% paraformaldehyde in 0.1M phosphate buffer (4% PFA). Brains were dissected with spinal cords attached and post-fixed in 4% PFA overnight. We discovered that tdT fluorescence can be easily visualized in intact brains and spinal cords by fluorescence epi-illumination. For this, brains with attached spinal cords are placed temporarily on a microscope slide and illuminated by epifluorescence (NG cube) using the 2X objective of an Olympus AX80 microscope. Images were taken at 2X and tiled to produce a complete low power reconstruction of tdT fluorescence in the spinal cord and brain. Subsequently, brains and spinal cords were returned to PBS prior to preparation either for light sheet microscopy (n=6, see below) or sectioning for immunocytochemistry (all other cases).

For sectioning, brains were cryoprotected in 27% sucrose, embedded in TissueTek O.C.T. (VWR International), and frozen. Blocks were sectioned by cryostat in the coronal plane at 30μm, and free-floating sections were collected in PBS at 480μm intervals through the cortex. One set of sections was mounted in serial order on microscope slides for visualization of native tdT fluorescence. Other sets of sections were immunostained for tdT.

### Assessment of co-labeling for tdT and GFP

In mice that received injections of retro-AAV/Cre at L2 and retro-AAV/GFP at C5, we assessed co-labeling of cortical neurons by immunostaining sections for GFP (FITC) and visualizing native tdT fluorescence in the same sections. For this, it was important to identify a GFP antibody that did not cross-react with the closely-related tdT protein. To test for cross-reactivity, sections from mice that received AAV/Cre only were immunostained using three commercial GFP antibodies (Invitrogen, rabbit anti-GFP, catalog #A11122, Abcam, goat anti-GFP, catalog #AB6673, and Novus, rabbit anti-GFP, catalog #NB600-308). All GFP antibodies were used at a dilution of 1-1000.

### Tissue clearing and light sheet fluorescence microscopy

Six brains were processed for light sheet microscopy (2 with C5 injections and 4 with L2 injections. Brains and attached spinal cords were immunostained and made optically transparent using a modified iDISCO+ whole-mount staining protocol (http://www.idisco.info); Renier et al., 2016). The following modifications were made to the iDISCO+ protocol: 1) The incubation step in permeabilization solution was followed by 2 additional 1-hour washes with Ptx.2. Tissue was incubated in 6-mL 1:600 (10 μl) Rabbit anti-RFP (Rockland, Lot# 35868) for 6 days. Secondary antibody amplification was done in 6-mL 1:600 (10 μl) Alexa Fluor 568 donkey anti-rabbit IgG (Thermo Fisher, Lot# 2044343) for 6 days.

Cleared brains were imaged on a Z.1 LSFM (Carl Zeiss Microscopy) modified for large, cleared samples in high refractive index solutions (Mesoscale Imaging System; Translucence Biosystems) using a 5×/0.16NA detection objective. Samples were mounted on a custom sample holder providing a sagittal view to the detection objective and submerged in dibenzyl ether (DBE). Each sample was illuminated through the left optic using a 561 nm laser line paired with a BP 575-615 emission filter at 60% power and 100 msec exposure. To construct a rendering of the whole sample, multiple z-stack scans were imaged in tiles with 20% overlap. Tiles were then stitched together in Vision4D (Arivis AG) and the completed image was rendered in Imaris 9.5 (Oxford Instruments) for analysis.

### Injections of biotinylated dextral amine (BDA) for orthograde tracing of CST projections

For intracranial BDA injections, mice were anesthetized with isofluorane, and positioned in a stereotaxic device. The scalp was incised at the midline and burr holes were placed in the skull overlying the right sensorimotor cortex. Injections were delivered as described below, and after the completion of the injections, the scalp was sutured and mice were placed on a water circulating heating pad at 37°C until they regained consciousness.

Table I summarizes the details on intra-cortical injections of BDA to trace CST projections from different cortical regions. As noted, some mice received mini-ruby conjugated BDA; the remainder received non-conjugated BDA. Both types of BDA were dissolved in 0.9% saline at a concentration of 10% weight/volume. Injections were made using either a Hamilton microsyringe fitted with a pulled glass micropipette or with a with an electronically-controlled injection system (Nanoliter 2000 injector and Micro4 pump controller, World Precision Instruments). Injections were at a depth of 0.5mm, and each individual injection was 0.4μl delivered over the course of 3-4 minutes.

Mice were allowed to survive for 14 days post-injection. Mice were euthanized with Euthasol® and perfused transcardially with 4% paraformaldehyde (PFA) in 0.1M phosphate buffer. Brains and spinal cords were dissected and stored in 4% PFA, and were transferred to 27% sucrose at 4°C for cryoprotection one day before being frozen for sectioning

For sectioning, the entire spinal cord was embedded in OCT, frozen, and cross-sections were taken throughout the rostro-caudal extent of the spinal cord from the spino-medullary junction through the sacral region maintaining serial order. Sections taken every 400μm were stained for BDA.

#### BDA staining

One set of floating sections through the spinal cord was stained with avidin-biotin peroxidase; another was stained for Cy3 fluorescence. For BDA staining with avidin-biotin-peroxidase, sections were washed in PBS with 0.1% Triton X-100 three times then incubated overnight in avidin-biotin-peroxidase complex (Vector Laboratories, PK-6100) in PBS with 0.1% Triton X-100 at room temperature. The following day, sections were washed in PBS 3X, then stained in nickel enhanced diaminobenzidine solution (Vector Laboratories, SK-4100) for 25 minutes. Sections were washed in PBS then mounted on gelatin-subbed slides, air dried, dehydrated, cleared, then coverslipped.

For BDA staining with Cy3 amplification, sections were washed in PBS, and then incubated in 1:200 dilution streptavidin-HRP (Perkin Elmer NEL75000-1ea) in PBS for 2 hours, washed 3 times in PBS, then reacted with TSA-Cy3 (1:100) in the supplied amplification diluent (Perkin Elmer SAT704A001ea). Sections were washed 2 times in PBS, then mounted on gelatin subbed slides, and coverslipped with Vectashield® (Vector labs H-1000).

### Quantification of CST axons at different spinal levels

Spinal cord cross-sections taken at 400μm intervals that were stained for BDA using avidin-biotin peroxidase were mounted in serial order on a single microscope slide. In each section, images were taken at 60X of BDA-labeled axons in the dorsal column (dorsal CST) and dorsal part of the lateral column (dorso-lateral CST). Images were imported into Photoshop, and BDA-labeled axons in each tract were counted, marking each as it was counted to avoid double-counting. Counts were assembled into a Prism file and displayed as graphs of numbers of BDA-labeled CST axons at different rostro-caudal locations.

## RESULTS

### Distribution of the cells of origin of corticospinal projections

Our initial approach to map the cells of origin of CST projections was done before the development of retro-AAV technology and used retrograde labeling with fluorogold (FG). The goal was to aid in defining coordinates to target populations of CST neurons within and outside of the canonical sensorimotor cortex. In mice that received bilateral injections of FG at C5, retrogradely label neurons were evident in the sensorimotor cortex, dorso-medial frontal cortex, and lateral cortex in the area of the S1 barrel field, in a pattern similar to what is seen in rats (Nielson et al 2011). Figure 1 illustrates the case with the largest number of retrogradely-labeled neurons.

**Figure 1.**
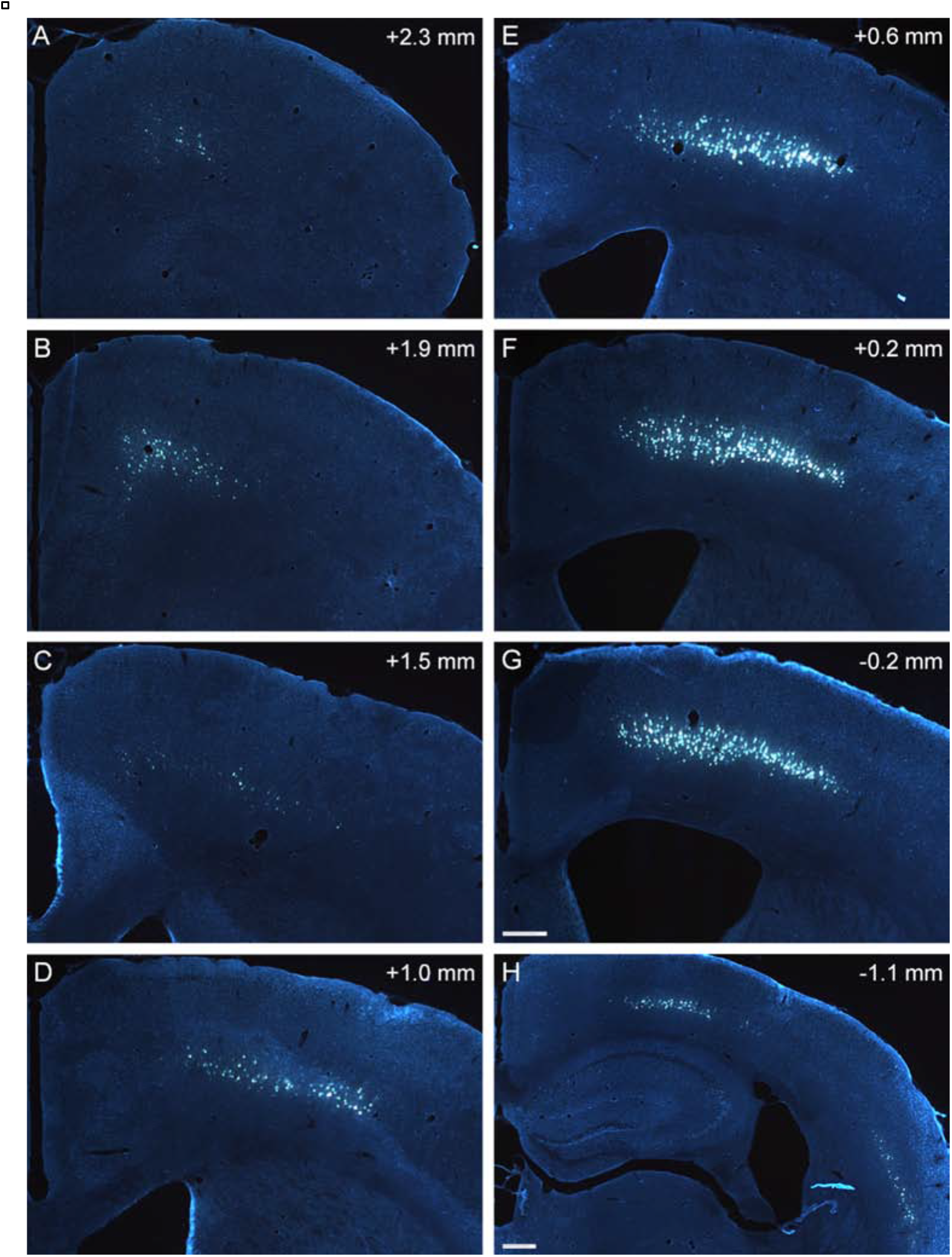
Rostrocaudal distribution of fluorogold (FG) labeled cell bodies in the cortex following injections of FG at C5. The panels illustrate the portion of the cortex containing retrogradely labeled neurons at different rostro-caudal levels. Coordinates with respect to bregma are indicated in the upper right of each panel. Scale bars = 400 μm in G (applies to A-G); 400μm in H.

FG-positive cortical motoneurons (CMNs) were evident in layer V throughout the primary motor, secondary motor and primary somatosensory cortical areas from approximately 2.3mm rostral to 1.5mm caudal to bregma (Fig. 1). We refer to this hereafter as the “main sensorimotor cortex”. Retrogradely-labeled cells were also present in the dorso-medial frontal cortex (centered at about 2.8mm anterior to bregma) in a region ~750 μm lateral to midline and ~600 μm deep. This area has been defined in rats through microstimulation studies as the “rostral forelimb area” (RFA) (Neafsey & Sievert 1982). The FG-labeled neurons in this rostral area are of approximately the same size and shape as the FG-labeled neurons in layer V in the main sensorimotor cortex. As in rats, there was also a spatially distinct population of FG-labeled cells in the lateral cortex posterior to bregma which also had the typical pyramidal form characteristic of the FG-labeled neurons in layer V of the main sensorimotor cortex (Fig.1I). This area was well-separated from the main sensorimotor cortex by an area with few if any retrogradely labeled neurons.

The cytoarchitecture of the two areas outside the main sensorimotor cortex that contained retrogradely-labeled neurons is illustrated in Figure 2. Each row illustrates the region containing retrogradely-labeled neurons in a section stained for cresyl violet and a nearby section immunostained for FG. Coronal sections through the dorso-medial frontal cortex are not perpendicular to the cortical surface, so cortical lamination is obscured. The area containing FG-labeled neurons in the dorso-lateral cortex is part of the S1 barrel field (Fig. 2G-I).

**Figure 2.**
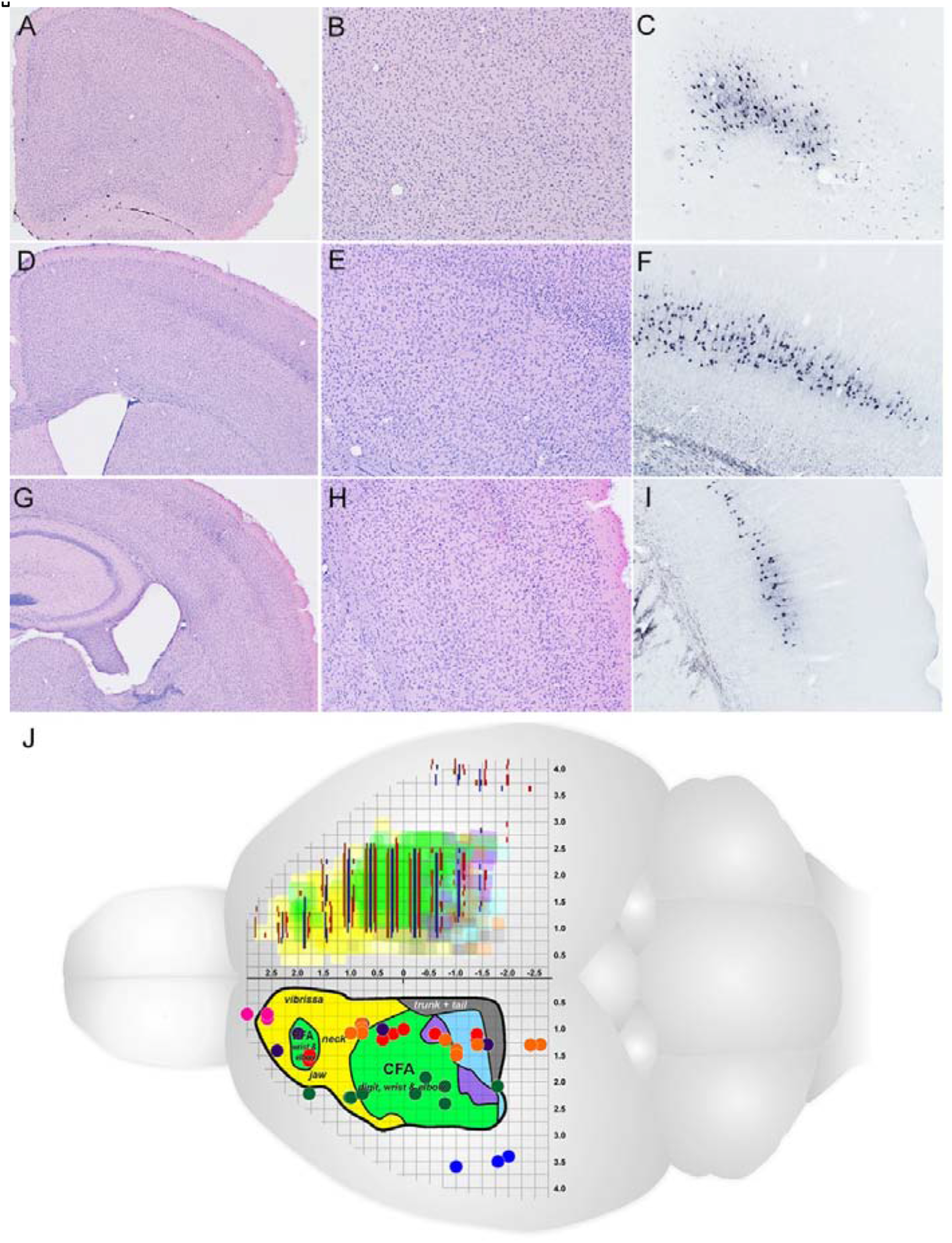
Map of retrogradely labeled cells and BDA injection sites. The different pairs of colored lines in the upper part of the figure illustrate the distribution of retrogradely labeled neurons from 3 mice. Each color corresponds to an individual mouse. Each pair of shaded colors represents the span of labeled cells in on the two sides of one specimen. The different colored dots in the lower part represent the location of BDA injection sites in the present experiment. Different colored dots represent different sets of targeted coordinates for injections on a particular date. Numbers indicate injection site locations for the respective figures that show the pattern of labeling of CST axons in the spinal cord resulting from injections at the indicated site. Some injection sites were plotted with slight offset to enable visualization of overlapping sites. Note the overlap with the rostral and caudal forelimb areas (RFA and CFA, respectively). J was created using a base drawing provided by Dr. T. Jones, which is a modified version of Figure 2B in Tennant et al., 2010.

To map the rostro-caudal distribution of retrogradely-labeled cells, we identified the location of bregma based on the appearance of the sections compared to the Paxinos mouse brain atlas (Paxinos 2004). We then created a summary diagram of the distribution of FG labeled neurons in the 3 cases by measuring the medio-lateral distance over which labeled cells were found in each section and superimposed this map on a microstimulation map of cortical motor function from (Tennant et al 2011). The medio-lateral distribution of cells at different rostro-caudal locations in each mouse is represented by the different colored lines over the right hemisphere of Figure 2J. Overall, the area occupied by retrogradely-labeled neurons overlapped with but was somewhat smaller than the motor map defined by micro-stimulation. The exception was the population of neurons in the area of the S1 barrel field, which is not represented in the micro-stimulation map.

### Retro-AAV transfection of CST neurons in tdT reporter mice

Retrograde labeling via retro-AAV/Cre in tdT reporter mice has several advantages over retrograde tracing by FG. First, genetically encoded fluorescent proteins extend throughout the dendritic arbor of neurons and also into axons. Second, whereas FG might be taken up by axons that pass through an injection site (Dado et al 1990), it has been reported that there is minimal uptake of retro-AAV by CST axons passing through the area of a spinal cord injection (Wang et al 2018). Third, labeling produced by retrograde expression of fluorescent proteins enables imaging by light sheet microscopy, allowing global visualization of the entire population of cortical neurons that project to the spinal cord (Wang et al 2018).

Here, we made intra-spinal injections of retrograde-AAV/Cre in transgenic Rosa^tdTomato^ and *Pten*^*loxP/loxP*^ /Rosa^tdTomato^ mice. Retrograde transfection with Cre induces bi-allelic expression of tdT leading to robust labeling of cell bodies, dendrites and axons. Remarkably, tdT fluorescence can be visualized in intact spinal cords and brains by fluorescence epi-illumination without clearing (Fig. 3A). This provides a convenient way to document the injection site and the cloud of fluorescently labeled cortical neurons prior to sectioning or preparation for iDISCO.

**Figure 3:**
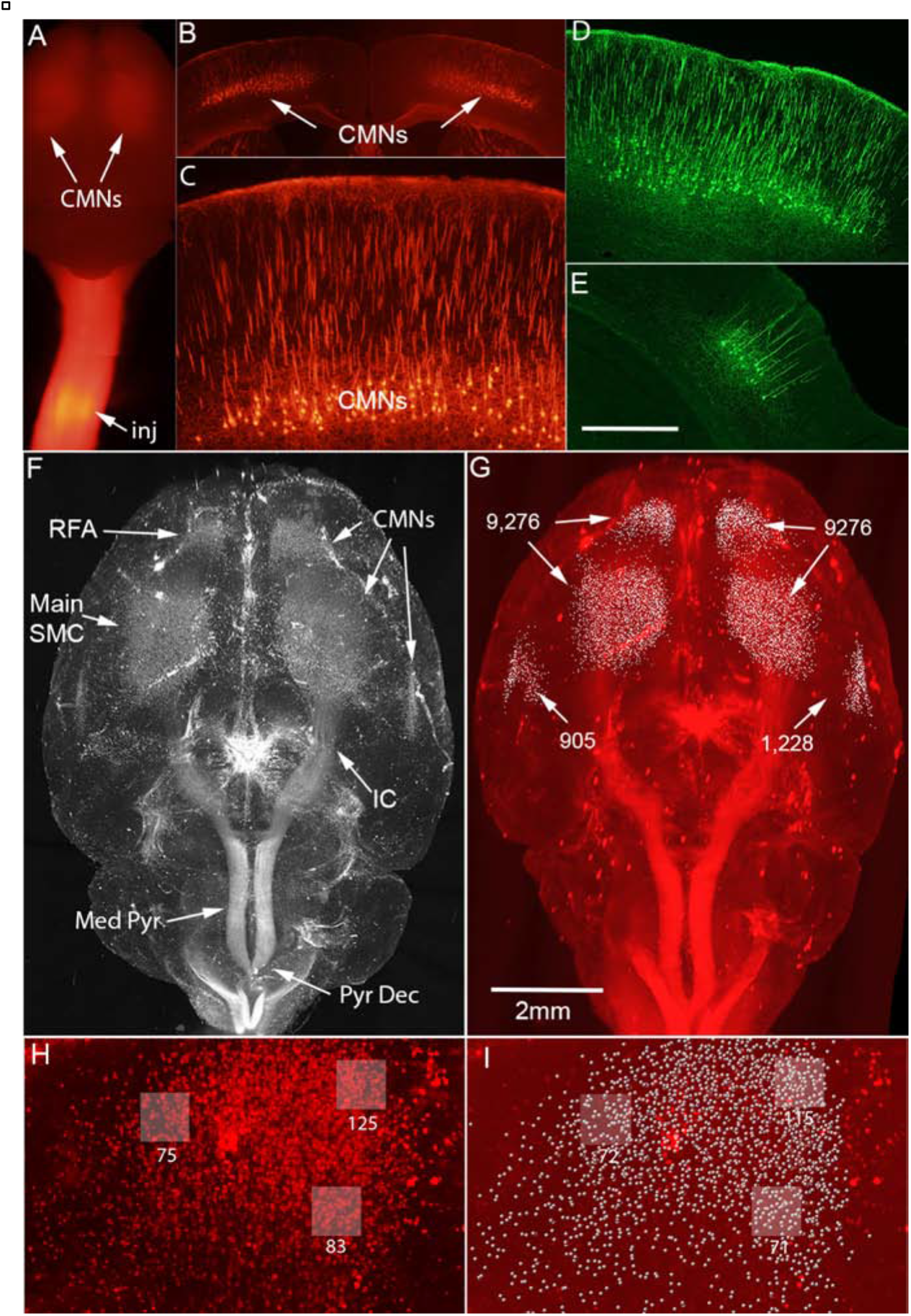
Retrograde transfection of cortical neurons following injections of retrograde-AAV/Cre at C5. A) Visualization of tdT fluorescence in intact un-cleared spinal cord and brain by fluorescence epi-illumination. B) Coronal section illustrating native tdT fluorescence in cell bodies and dendrites of pyramidal neurons in layer V. C) Higher magnification view reveals intense tdT fluorescence in nuclei and labeling throughout dendrites including fine dendritic arbors in layer I-II. D) Immunostained section (FITC) illustrating tdT-positive neurons in the main sensorimotor cortex. E) Immunostained section illustrating tdT-positive neurons in the lateral cortex near the barrel field. F) Projection image of 3D reconstruction with light sheet microscopy after clearing and immunostaining with iDISCO. Thousands of tdT-positive cortical motoneurons (CMNs) are evident in layer V throughout the primary motor, secondary motor and primary somatosensory cortical areas and the lateral cortex near the barrel field. IC=internal capsule; Med Pyr=medullary pyramid; Pyr Dec=pyramidal decussation. G) Labeled neurons identified by “Spots” function in Imaris. Numbers of labeled neurons in the main sensorimotor cortex and lateral cortex are indicated.

In coronal sections through the brain, native tdT fluorescence can be seen without immunostaining in cell bodies and dendrites of pyramidal neurons in layer V including fine dendritic arbors in layer I-II (Fig. 3B&C). Of note, native tdT fluorescence was most intense in nuclei. As with retrograde labeling with FG, tdT-positive neurons were present in the main sensorimotor cortex, the dorso-medial frontal cortex (rostral forebrain area, RFA), and the lateral cortex near the barrel field. With immunostaining, tdT labeling extended throughout the dendritic arbor revealing a dense network of dendrites in layers V and I/II (Figure 3D). Figure 3D illustrates the main sensorimotor cortex and Fig. 3E illustrates labeled neurons in the lateral cortex near the barrel field. In both sites labeled CST neurons had a similar morphology, with a large pyramidal-shaped cell body, a single thick apical dendrite extending toward the cortical surface and extensive branches in layers I&II and multiple basal dendrites in the deep part of layer V.

3D imaging of a different mouse with light sheet microscopy after clearing and immunostaining with iDISCO revealed thousands of tdT-positive CST neurons in layer V throughout the primary motor, secondary motor and primary somatosensory cortical areas (Fig. 3D and movie 0315G). Clusters of tdT-positive neurons in the dorso-medial frontal cortex and the lateral cortex near the barrel field were separated from the main sensorimotor cortex by an area with few labeled neurons. Descending axons of the CST were prominently labeled in the 3D light sheet renderings in the subcortical white matter and internal capsule, medullary pyramid, and pyramidal decussation (Fig. 3D and movie 0315G).

We used the Imaris “Spots” function to estimate the number of retrogradely-labeled CST neurons in the 3-Dimensional light sheet reconstructions. First, the selection box was positioned to include the area containing labeled neurons in the main sensorimotor cortex on one side. Next, tdT positive neuronal cell bodies were segmented using the “quality” thresholding procedure to recognize labeled neuronal cell bodies but exclude artifacts and labeled dendrites. Then, artifactual hits at the surface of the brain, along blood vessels, and outside the area containing retrogradely labeled neurons were manually edited. The process was repeated on the opposite side and then for the two areas in the dorso-lateral cortex with tdT positive neurons. Figure 3G illustrates the above threshold hits in the 4 sampled areas with retro-AAV/Cre injections at C5. Before editing, there were 10,883, above threshold hits in the main sensorimotor cortex on the left side and 10,874 on the right. After excluding non-neuronal labeling and scattered hits outside the sensorimotor cortex, there were 9,276 labeled CST neurons on each side. Counts in the lateral cortex near the barrel field revealed 1,534 hits on the left side and 1,437 on the right before editing and 905 and 1,228 labeled neurons after editing for an average of 1067 labeled CST neurons per side.

To assess the extent to which the “Spots” quality thresholding included artifacts and/or failed to detect tdT-positive neuronal cell bodies, we compared “Spots” counts with manual counts in sample areas with a moderate number of labeled neurons (the RFA on the left). Manual counts in three 150X150μm areas (shaded boxes in Figure 3H) yielded counts of 75,125, and 83 tdT-positive neurons and the corresponding “Spots” fields had 72, 115 and 71 (Figure 3I), which represents a difference of 4%, 8% and 14% (average of 8.7%). Large bright fluorescent artifacts were not recognized. Similarly, close inspection in the area of dense retrograde labeling in the main sensorimotor cortex indicates that the majority of labeled neurons were correctly recognized and few few artifacts were incorrectly recognized as cells.

With injections of retro-AAV/Cre into lumbar regions (L2 vertebral level), retrogradely transfected TdT-positive layer V neurons were distributed in a smaller region of the sensorimotor cortex extending from about bregma to 1.8mm posterior and from 0.8-1.8mm lateral to the midline (Figure 4 and movie 0612A). Figure 4A-D illustrates a series of sections at 0.5mm intervals from one mouse and Figure 4E illustrates a light sheet reconstruction from a different mouse. Of note, when viewed from above (as in Fig. 4E), the dense network of labeled dendrites in layer I/II creates a haze-like cloud over the labeled neurons in layer V. No labeled neurons were seen in the lateral cortex in the region of the barrel field in mice with L2 injections. Counts using Imaris (Fig. 4F) revealed 1,394 above threshold hits on the left side and 1,517 on the right before editing. After excluding non-neuronal labeling and scattered hits outside the sensorimotor cortex, there were 1,161 labeled CST neurons on the left and 1,218 on the right for an average of 1,770 labeled CST neurons per side.

**Figure 4:**
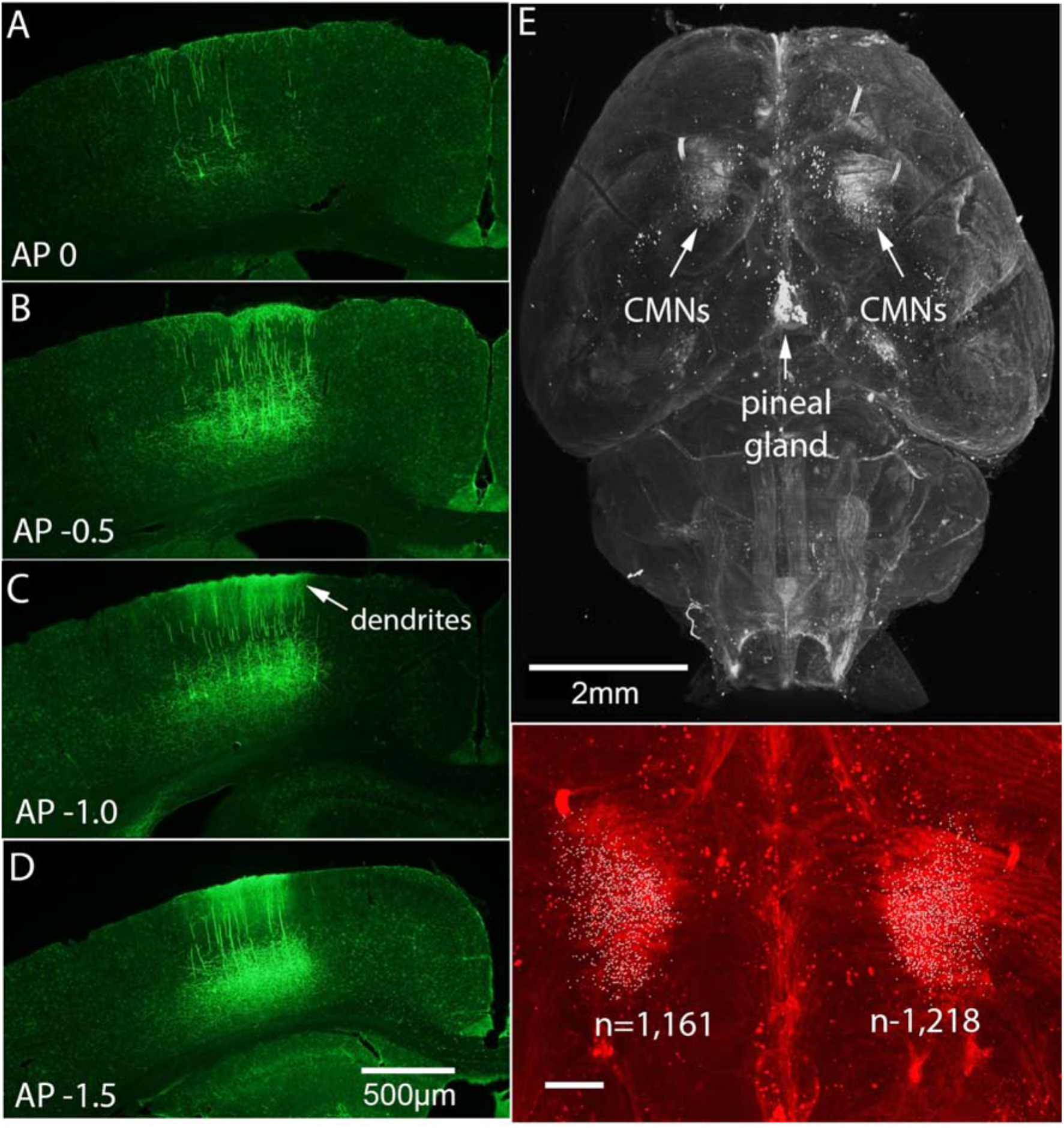
Retrograde transfection of cortical neurons following injections of retrograde-AAV/Cre at L2. A-D) Immunostained sections (FITC) illustrating tdT-positive neurons in the main sensorimotor cortex at different locations with respect to bregma. E) Projection image of 3D reconstruction with light sheet microscopy after clearing and immunostaining with iDISCO. TdT-positive cortical motoneurons (CMNs) are present in posterior sensorimotor cortex. F) Labeled neurons identified by “Spots” function in Imaris. Numbers of labeled neurons in the main sensorimotor cortex are indicated. Scale bar in F=500μm.

Comparison of the distribution of labeled neurons with injections at C5 vs. L2 suggests possible overlap in areas posterior to bregma. To determine the extent of overlap of CST neurons labeled by C5 vs. L2 injections, we injected retro-AAV/Cre at L2 and retro-AAV/GFP at C5, immunostained brain sections for GFP (FITC), and then imaged sections for GFP immunofluorescence and native tdT fluorescence. For this, it was important to identify a GFP antibody that did not cross-react with the closely-related tdT protein. To test for cross-reactivity, sections from mice that received AAV/Cre only were immunostained using three commercial GFP antibodies (Invitrogen, rabbit anti-GFP, catalog #A11122; Abcam, goat anti-GFP, catalog #AB6673; and Novus, rabbit anti-GFP, catalog #NB600-308). All GFP antibodies were used at a dilution of 1-1000. As illustrated in Figure 5, there was robust immunostaining of tdT-positive CST neurons when sections were immunostained using GFP antibodies from Abcam and Invitrogen. However, there was no detectable immunofluorescence when sections were immunostained using the GFP antibody from Novus. Based on these results, we used the Novus antibody for assessment of double-labeling.

**Figure 5:**
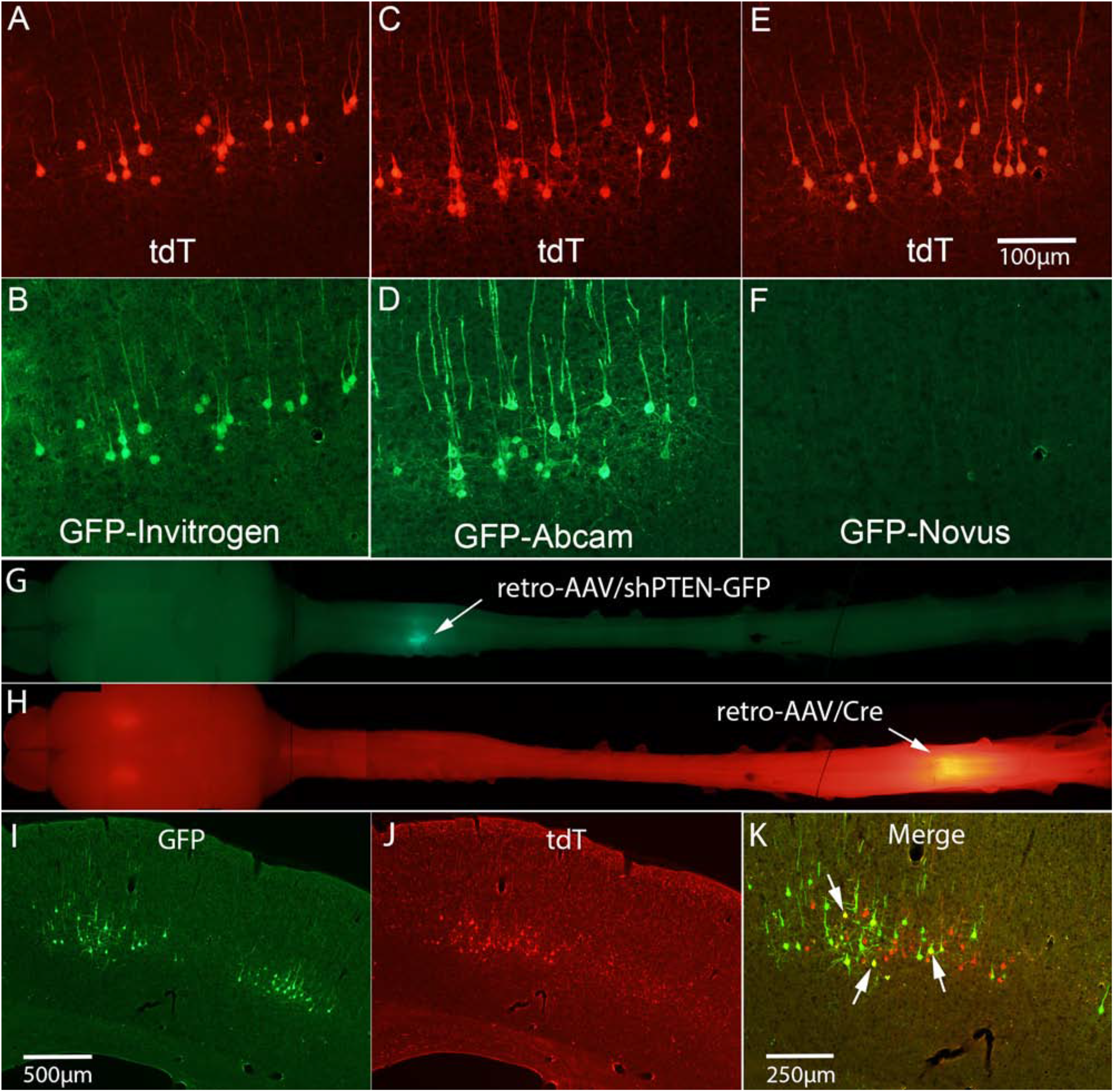
Retrogradely labeled CST neurons with retro-AAV/Cre injections at L2 and retro-AAV/shPTEN-GFP at C5. A-E: Some commercial GFP antibodies recognize tdT. A) Native tdT fluorescence in a section stained for GFP using Invitrogen antibody. B)Immunofluorescence for GFP in same section shown in A. C) Native tdT fluorescence in a section stained for GFP using Abcam antibody. D)Immunofluorescence for GFP in same section shown in C. E) Native tdT fluorescence in a section stained for GFP using Novus antibody. F) Immunofluorescence for GFP in same section shown in E. G) Visualization of GFP fluorescence in intact un-cleared brain and spinal cord by fluorescence epi-illumination. H) Visualization of tdT fluorescence in the same brain and spinal cord. I) Coronal section through posterior sensorimotor cortex approximately 1.8mm caudal to bregma. J) Same section illuminated to reveal native tdT fluorescence. K) Merged image of C and D to reveal the few double-labeled neurons that were present (arrows). Scale bar in C=500μm and applies to C&D; scale bar in E=250μm.

The distribution of retrogradely transfected GFP-positive neurons was similar to that reported above with retro-AAV/Cre injections at C5. In the caudal part of the region containing GFP-labeled neurons (approximately 1mm caudal to bregma), GFP-positive neurons were found in two mediolateral locations separated by a space with few if any GFP-positive neurons (Fig. 5A). TdT-positive neurons labeled by AAV/Cre injections at L2 were detected in the same sections by native fluorescence of tdT (no IHC); These were present in the space between the populations of GFP-positive CST neurons, overlapping the medial area containing GFP-positive neurons (Fig. 5B). Although there was overlap in the distribution of GFP- and tdT-positive CST neurons, there were only a few double-labeled neurons. Double-labeled neurons may represent neurons that project to both cervical and lumbar levels or may indicate that injections of AAV/Cre at C5 can lead to transfection of some CST axons passing through cervical levels *en route* to lower spinal levels. In either case, the number of double-labeled neurons is small.

Although the focus here is on the cells of origin of the CST, cells of origin of other descending pathways to the spinal cord are also labeled following AAV/Cre injections at C5 and L2 including neurons in the red nucleus and reticular formation, vestibular nucleus, and hypothalamus. These can be discerned in the light sheet movies and are evident in cross-sections through the brain (not shown). Of note, with injections at L2, labeled neurons were also seen in Barrington’s nucleus which gives rise to descending pathways that are important for bladder function (not shown).

One other aside is that the light sheet images reveal the presence of tdT in the pineal gland (see especially Figure 5E). We reported this phenomenon in abstract form with intra-cortical injections of AAV (Steward et al., 2016), and a manuscript documenting accumulation of AAV in the pineal gland after injection in different CNS sites is in preparation.

### BDA-tracing of corticospinal projections from different cortical areas

To assess projections from different populations of CMNs, we made single injections of BDA at different locations. Injection sites are mapped on Figure 2J; the center of the tracer injection in each mouse is indicated by a colored dot and different colors indicate sets of mice with injections targeted to particular coordinates (see Table 1).

### Rostral part of the main sensorimotor cortex (forelimb representation)

With BDA injections into the rostral part of the main sensorimotor cortex, labeled CST axons were restricted primarily to cervical levels. In three mice (1120A-C), BDA injections were centered at 1.8mm anterior, and 1.5-1.6 lateral (medial part of the forelimb cortex; M1). Figure 6 illustrates the case (1120C) with the largest number of labeled axons. The injection site is illustrated in Fig. 6G and the cytoarchitecture of the area is illustrated in a nearby cresyl violet stained section in Fig. 6H. The rostro-caudal distribution of CST axons is illustrated in Figure 6A-E. The number of labeled axons in all tracts decreased rapidly moving from rostral to caudal (Fig. 6A-E); few labeled axons extended to the mid-thoracic level (Fig. 6E), and none extended to lower thoracic levels.

**Figure 6:**
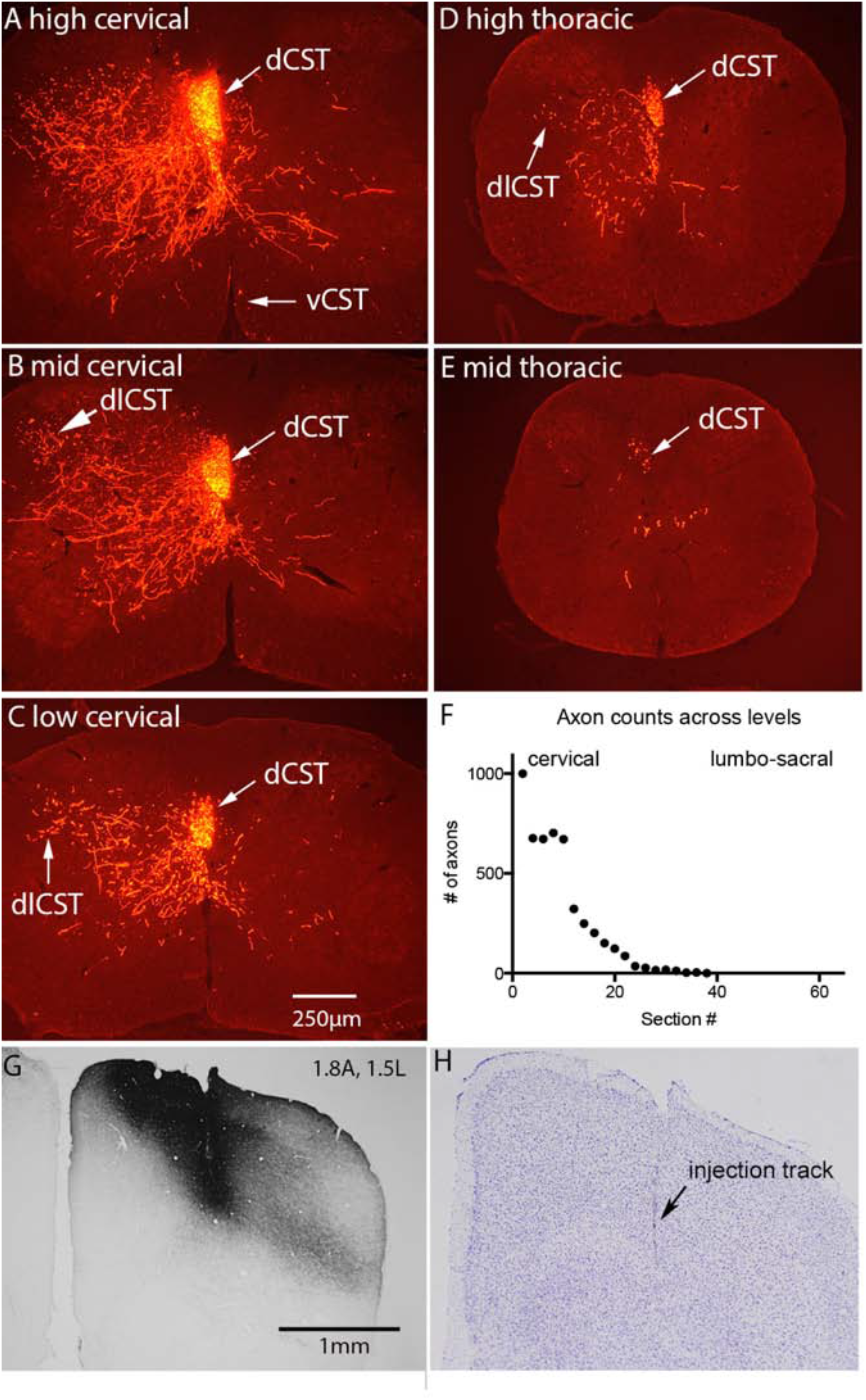
Antero-lateral forelimb cortex projects selectively to cervical and high thoracic levels with terminal arbors located throughout the intermediate lamina and medial ventral horn. The panels illustrate Cy3-labeled axon arbors at cervical-high thoracic levels following a single injection of BDA at 1.8A, 1.5L. A) high cervical level; B) mid cervical; C) low cervical; D) high thoracic; E) mid thoracic; F) Counts of labeled axons in the dCST in sections spaced at 800μm intervals; G) BDA labeling at injection site; H) Cresyl violet stained section at injection site. Abbreviations: dCST=dorsal corticospinal tract in the dorsal column; dlCST=dorsolateral corticospinal tract in the dorsal part of the lateral column; vCST=labeled axons in the ventral column ipsilateral to the cortex of origin in the expected location of the ventral CST.

To quantify the rostro-caudal extension of BDA-labeled CST axons, we counted labeled axons in the dCST in sections spaced at 800μm intervals (Fig. 6F). In the first 5 sections in the rostro-caudal series, counts of labeled CST axons in the dCST ranged from 672 to over 1000 (average of 773). The number of labeled axons decreased progressively moving caudally, and no labeled axons extended further than mid-thoracic levels (section #38, which is 14.4mm from the rostral end of the block). In addition to the main tract in the dlCST, there were 65 labeled axons in the dlCST at mid cervical levels (Fig. 6B) and 30 labeled axons in the dCST ipsilateral to the injection. In this case, there were also 7 BDA labeled in the ventral column contralateral to the main labeled tract in the position expected for the vCST (arrow, Fig. 6A). Labeled axons in the vCST were present in only a minority of mice as noted below.

At cervical levels, labeled axons extended from the dCST and dlCST into the gray matter, where they gave rise to dense terminal arbors in the medial parts of laminae 4-7 (Fig. 6A, B&D). Few labeled arbors extended into the lateral part of the ventral horn (Fig. 6A-C). Labeled axons extended into the dorsal horn, but not into lamina 1-3. A moderate number of axons extended across the midline to arborize in the gray matter of the medial ventral horn on the contralateral side (Fig. 6A-D). The density of the arbors decreased moving caudally in parallel with the number of axons in the dCST.

In cases with injections at 0.6-0.8A, 0.9L, most axons terminated at cervical levels but some extended through thoracic and a few extended to high lumbar levels. Fig. 7, illustrates case 0225H, where the injection was at 0.8A, 0.9L (Fig. 7G). Counts of labeled axons in the dCST at cervical levels ranged from 248-349 (average of 255 in the first 9 sections of the series), with 10 labeled axons in the dCST contralateral to the main labeled tract and no labeled axons in the ventral CST. About 10% of the labeled axons extended in the dCST into thoracic levels and a few extended into high lumbar segments (Fig. 7E; insets illustrate labeled axons in the dCST and labeled arbors in gray matter). Only one axon extended beyond section 44 (17.6mm from the rostral end of the block), and this single axon ended in section 52 (20.8mm from the rostral end of the block). At cervical levels, labeled terminal arbors extended throughout the medial half of lamina 5-7 with a few axons extending to the ventro-medial boundary of the ventral horn (Figure 7B). At mid-thoracic levels, axons could be seen streaming from the dCST into the medial-dorsal part of lamina 7. In contrast to the case illustrated in Fig. 6, there are very few re-crossing axons.

**Figure 7:**
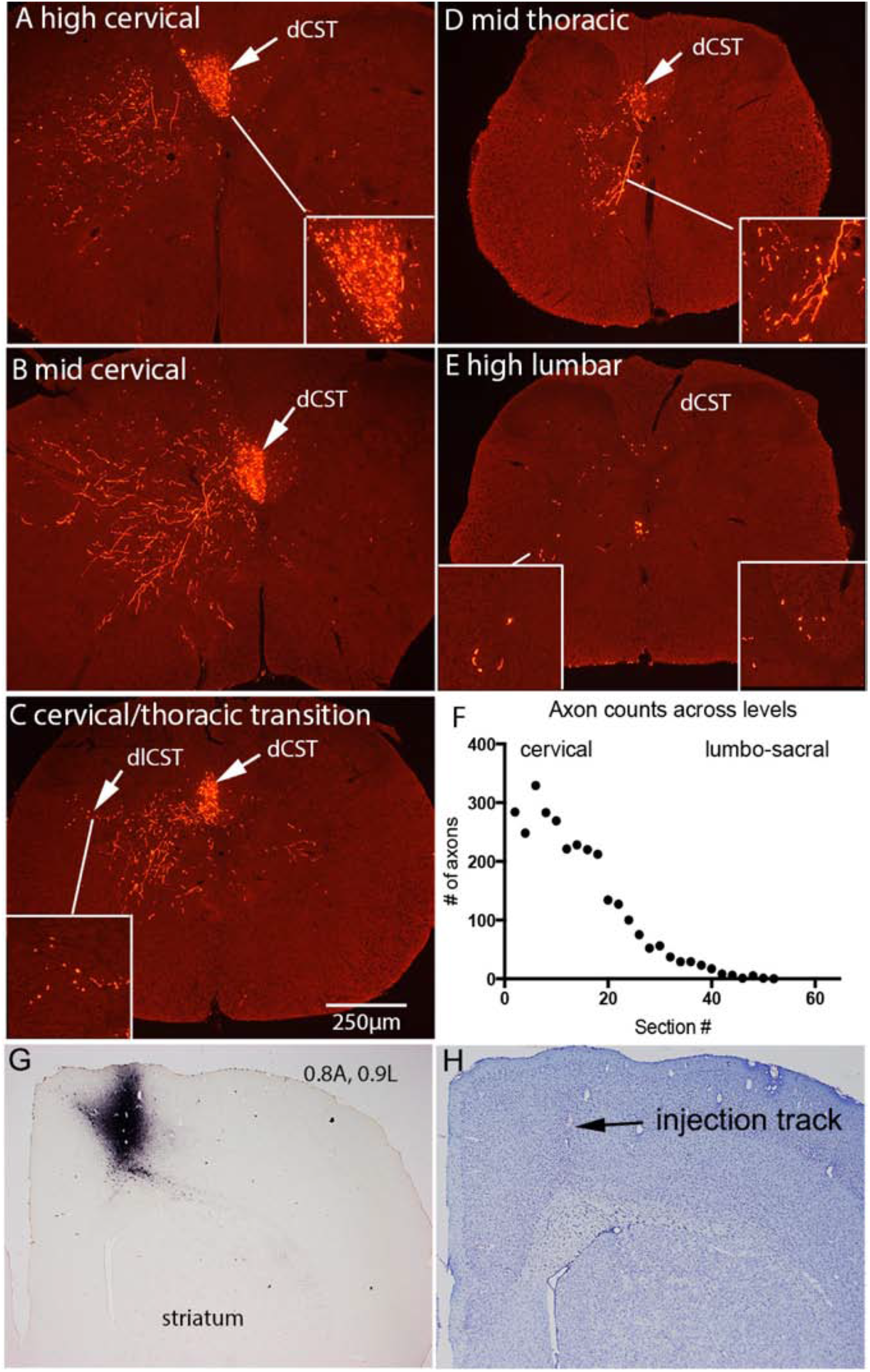
Medial posterior forelimb cortex projects selectively to cervical and mid thoracic levels with terminal arbors located throughout the intermediate lamina and medial ventral horn. The panels illustrate Cy3-labeled axon arbors at cervical-mid thoracic levels following a single injection of BDA at 0.8A, 0.9L. A) high cervical level; B) mid cervical; C) low cervical; D) high thoracic; E) mid thoracic; F) Counts of labeled axons in the dCST in sections spaced at 800μm intervals; G) BDA labeling at injection site; H) Cresyl violet stained section at injection site. Abbreviations are as in Fig. 6.

In 3 mice (1120D-F), injections were into the posterior part of the medial forelimb cortex (target coordinates for injections were at bregma and 1.0 lateral and the actual coordinates were 0, 0.2 and 0.4mm anterior and 1.0, 1.1, and 1.2mm lateral). In the case with the fewest labeled axons (about 80 BDA-labeled axons in the dorsal CST at high cervical levels), projections were to the cervical region only as in the cases injected at 0.5-1.0 anterior. In the two other cases, labeled axons extended to lumbar and sacral levels. Figure 8 illustrates the case with the largest number of labeled axons (1120E). The high density of labeled axons in the dCST contralateral to the injection precluded accurate counts, but there were 50 labeled axons in the dCST ipsilateral to the injection, 40 labeled axons in the dlCST, and 2 labeled axons in the vCST. Labeled axons in the dlCST are especially evident at thoracic levels (Fig. 8C). There were also 8 labeled axons in the lateral column contralateral to the main labeled tract (arrows with asterisk in Figure 8C); this is an unusual location for CST axons, but has been previously been reported in mice (Zheng et al 2006) and rats (Brosamle & Schwab 1997).

**Figure 8:**
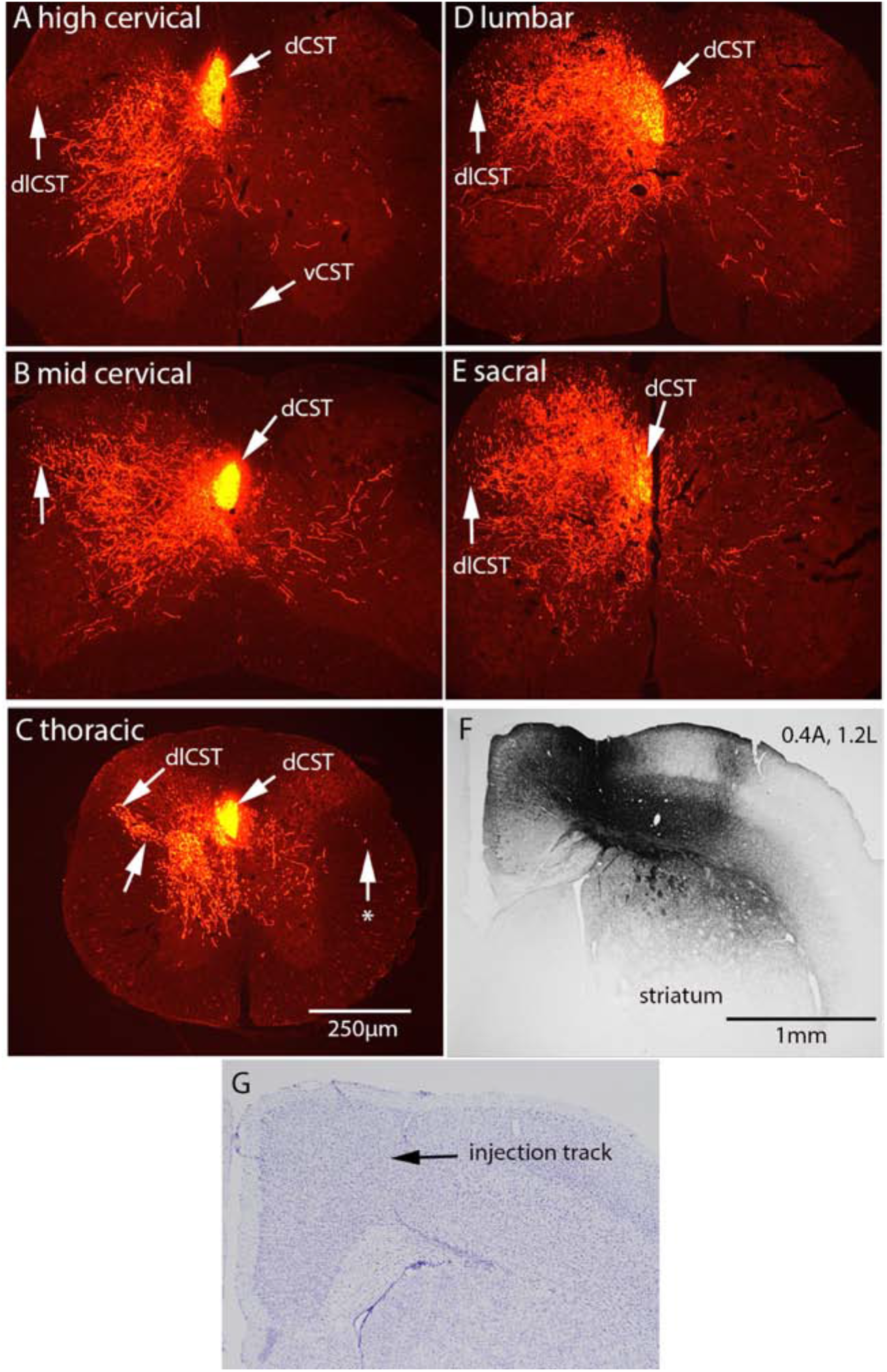
Medial posterior forelimb cortex/transition to anterior hindlimb cortex projects to cervical-sacral levels with terminal arbors throughout the intermediate lamina and medial ventral horn. The panels illustrate Cy3-labeled axon arbors at cervical-sacral levels following a single injection of BDA at 0.4A, 1.2L. A) high cervical level; B) mid cervical; C) low cervical; D) high thoracic; E) mid thoracic; F) BDA labeling at injection site; G) Cresyl violet stained section at injection site. Abbreviations are as in Fig. 6. Asterisk in C indicates a few labeled axons in the lateral column ipsilateral to the injected cortex.

Labeled axon arbors in the gray matter were dense at all rostro-caudal levels. At cervical levels (Fig. 8A&B) dense axon arbors were present in the medial part of the ventral horn (lamina 7) extending through lamina 4 of the dorsal horn but were largely absent from lamina 2-3. At thoracic levels, the highest density of labeled arbors was in lamina 5 (Figure 8C); of note, labeled axons could be seen streaming from the dlCST toward the dense arbors in lamina 5 (Fig. 8C, unlabeled arrow). There were also large numbers of labeled arbors in in the dorso-medial part of lamina 7 of the ventral horn. At lumbar and sacral levels, labeled arbors were dense in lamina 4-5 and lamina 7-8 of the ventral horn (Fig. 8D&E). At all levels, a moderate number of labeled axons re-crossed the midline to arborize on the side contralateral to the labeled tract.

### Lateral forelimb cortex

With BDA injections into the lateral part of the forelimb cortex, labeled axons also were also restricted primarily to cervical levels, but the distribution of labeled terminal arbors in the gray matter of the spinal cord was different.

Figure 9 illustrates a case (0812D) in which the BDA injection was centered in the anterior-lateral forelimb cortex 1mm anterior and 2.3mm lateral (lateral M1/S1 border). The injection site is illustrated in Fig. 9G and Fig. 9H illustrates a nearby section stained for cresyl violet. The rostro-caudal distribution of CST axons is illustrated in Figure 9A-E and counts of labeled axons across levels are illustrated in Fig. 9F. At high cervical levels, counts revealed an average of 510 labeled axons in the dCST contralateral to the injection in the first 6 sections of the series and 10 labeled axons in the dlCST (Fig. 9A-C). Of note, there were also a large number of labeled axons in the dCST ipsilateral to the injection (average of 199 in the first 6n sections of the series. The number of labeled axons in the dCST decreased sharply moving from cervical to thoracic levels. Very few axons extended to lumbar levels (5 labeled axons are present at the level indicated in Fig. 9E).

**Figure 9:**
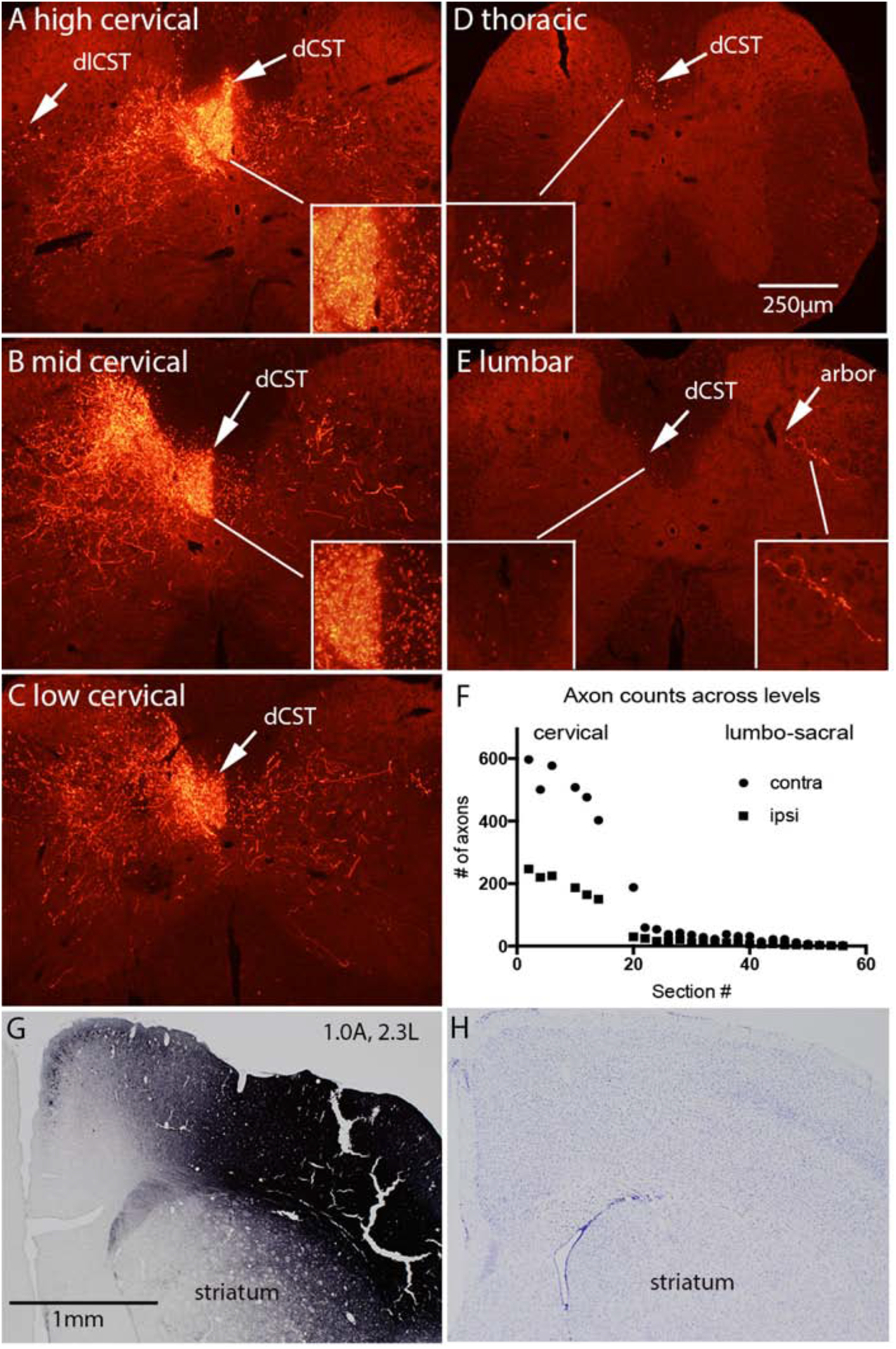
Lateral posterior forelimb cortex (bregma-1.0A) projects selectively to cervical levels with terminal arbors in the dorsal intermediate lamina and medial ventral horn. The panels illustrate Cy3-labeled axon arbors at cervical but not more caudal segments following a single injection of BDA at 1.0A, 2.3L. A) high cervical level; B) mid cervical; C) low cervical; D) high thoracic; E) mid thoracic; F) Counts of labeled axons in the dCST in sections spaced at 800μm intervals; G) BDA labeling at injection site; H) Cresyl violet stained section at injection site. Abbreviations are as in Fig. 6. Note large number of labeled axons in the dCST ipsilateral to the cortex of origin (insets in A and B) and that although very few axons extend to lumbar levels, there are occasional arbors in the gray matter (panel E, arbor).

In contrast to what was seen with injections into the medial forelimb cortex, BDA labeled arbors were most dense in the medial part of lamina 4-6, and fewer axons extended into lamina 7 of the ventral horn (Figure 9A-C). There were a moderate number of labeled axon arbors in laminae 4-6 on the contralateral side. The density of the arbors decreased moving caudally in parallel with the decrease in numbers of BDA-labeled axons in the dCST. Despite the low number of labeled axons that extended to lumbar levels, there were occasional labeled arbors in the gray matter (Fig. 9E illustrates an arbor on the side ipsilateral to the cortex of origin).

Figure 10 illustrates one of 3 cases in which the BDA injection was centered in the posterior-lateral forelimb cortex. In this case (0812I), the injection was centered at 0.8mm posterior 2.4mm lateral (Fig. 10G). The rostro-caudal distribution of CST axons is illustrated in Figure 10A-E and counts of labeled axons across levels are illustrated in Fig. 10F. In all 3 cases, there were several hundred BDA-labeled axons in the dorsal CST at high cervical levels (average of 360 labeled axons in the first 9 sections of the series in Figure 10F). There were a few labeled axons (4,4, and 5 in the 3 different mice) in the dCST contralateral to the main labeled tract and a moderate number of labeled axons (15,20, and 20 respectively) in the dlCST at cervical levels. One of the 3 mice had about 6 labeled axons in the vCST contralateral to the main labeled tract; the other 2 mice had none. The number of labeled axons in the dCST remained high through mid-thoracic levels (Figure 10C), dropping off at lumbar levels (Figure 10D). Only a few labeled axons continued to low lumbar levels (Figure 10E, inset).

**Figure 10:**
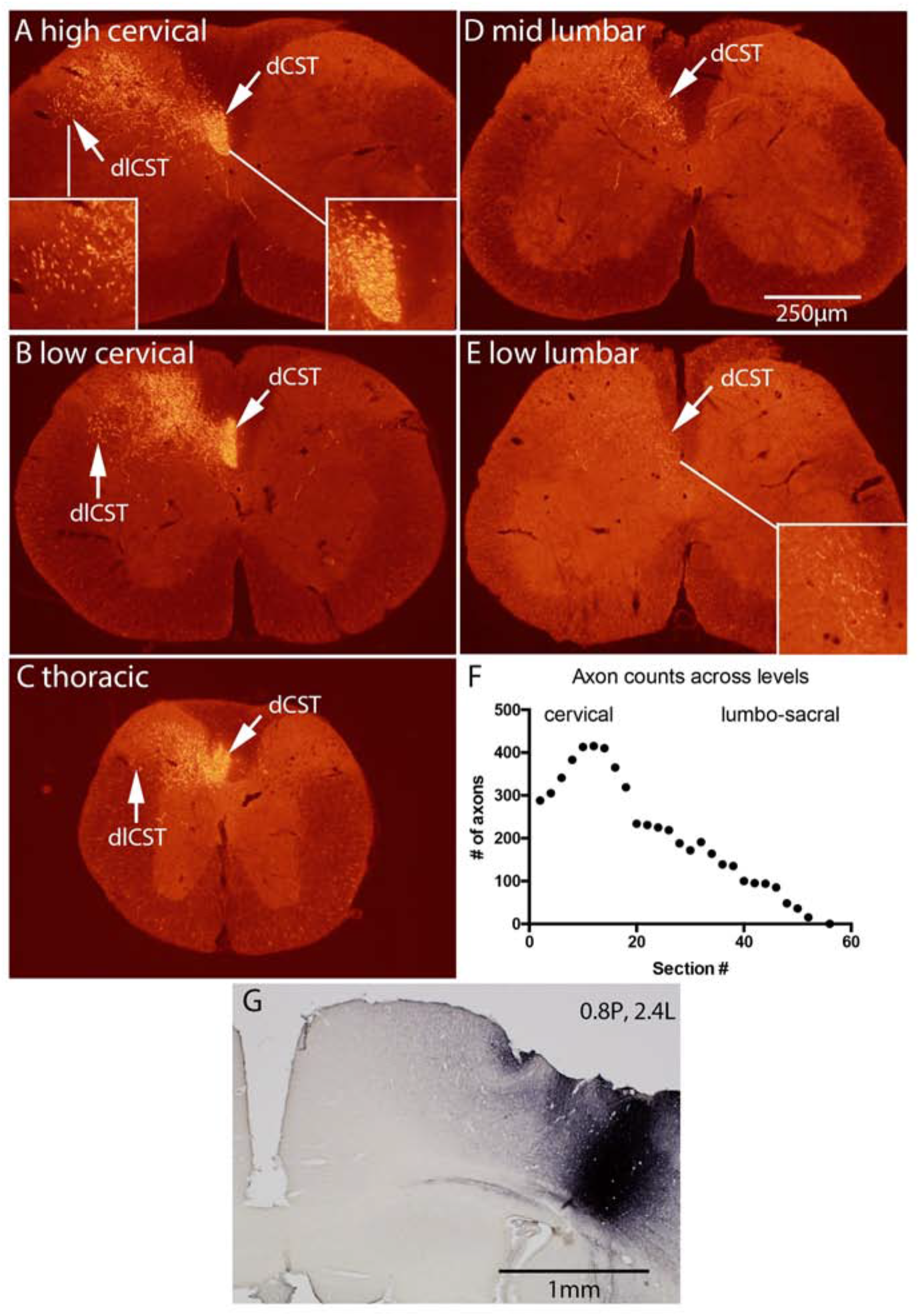
Lateral posterior forelimb cortex (1.0 posterior to bregma) projects to cervical-lumbar levels with terminal arbors in the deep layers of the dorsal horn. The panels illustrate Cy3-labeled axon arbors at cervical-lumbar following a single injection of BDA at 0.8P, 2.4L. A) high cervical level; B) mid cervical; C) low cervical; D) high thoracic; E) mid thoracic; F) Counts of labeled axons in the dCST in sections spaced at 800μm intervals; G) BDA labeling at injection site. Insets show high magnification views of labeled axons.

At high cervical levels, labeled arbors were concentrated in the intermediate lamina and middle part of lamina 3-4 in the dorsal horn. Very few axons extended into the ventral horn. At low cervical levels-thoracic levels, labeled arbors were in the medial part of lamina 3-5 of the dorsal horn and upper intermediate lamina. A noteworthy feature with injections in the posterior-lateral forelimb region site was the relative lack of re-crossing projections. In all 3 mice, there were elaborate arbors ipsilateral to the main labeled tract but very few axons extended across the midline to the side contralateral main labeled tract.

### Projections from the caudal sensorimotor cortex (hindlimb representation)

With injections into the caudal sensorimotor cortex, BDA-labeled axons extend through cervical levels without sending extensive collaterals into the gray matter and then arborize extensively at thoracic-sacral levels. Figure 11 illustrates a case (1120I) in which the BDA injection was centered 0.6mm posterior and 1.1mm lateral. The injection site is illustrated in Fig. 11G and Fig. 11H illustrates a nearby section stained for cresyl violet. The rostro-caudal distribution of CST axons is illustrated in Figure 11A-E and counts of labeled axons across levels are illustrated in Fig. 11F. Throughout cervical levels (first 10 sections of the series), counts revealed an average of 717 labeled axons in the dCST contralateral to the injection, with 25 labeled axons in the dCST contralateral to the cortex of origin, and 35 in the dlCST (Figure 11A-C; insets show high magnification views of the dlCST). There were no labeled axons in the ventral column ipsilateral to the injection in the position of the vCST. The number of labeled axons in the dCST remained high (average of 704) throughout thoracic and lumbar levels, and 178 axons extend to sacral levels (graph in Figure 11F).

**Figure 11:**
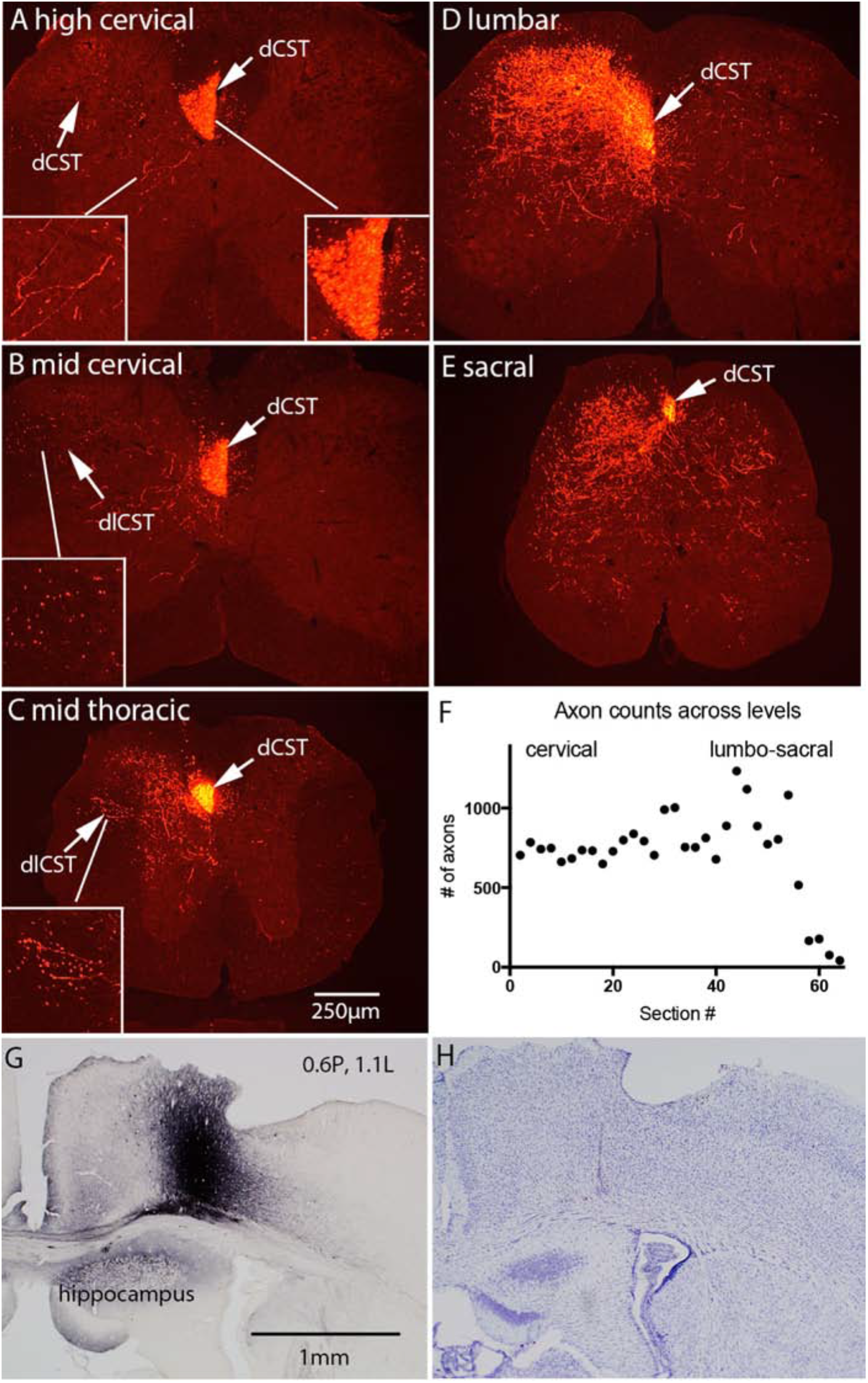
Medial forelimb cortex projects selectively to cervical levels with terminal arbors located primarily in the intermediate lamina and ventral horn. The panels illustrate Cy3-labeled axon arbors at cervical but not more caudal segments following a single injection of BDA at 0.6A, 1.1L. A) high cervical level; B) mid cervical; C) low cervical; D) high thoracic; E) mid thoracic; F) Counts of labeled axons in the dCST in sections spaced at 800μm intervals; G) BDA labeling at injection site; H) Cresyl violet stained section at injection site. Abbreviations are as in Fig. 6. Arbors extend into intermediate lamina and dorso-medial ventral horn. Insets show axons in the dlCST and dCST at 2X higher magnification.

There were very few BDA-labeled arbors in the gray matter at cervical levels (Fig. 11A&B). The density of arbors in the gray matter increased progressively moving caudally (Fig. 11C-E), reaching maximal density at lumbar levels (Fig. 11D). At thoracic levels, arbor density was highest in lamina 4-5 of the dorsal horn and a few axons extended into lamina 7 in the ventral horn (Figure 11C). At lumbar levels, very elaborate axon arbors extended throughout lamina 3-5 of the dorsal horn, and a few axons extended into lamina 7 in the ventral horn. Some labeled axons extended across the midline at lumbar levels, and there were large numbers of re-crossing axons in sacral segments (figure 11E).

Figure 12 illustrates a case (0321H) in which the BDA injection was centered in the caudal hindlimb region 2.0mm posterior and 1.5mm lateral (Fig. 12G). The rostro-caudal distribution of CST axons is illustrated in Figure 12A-E and counts of labeled axons across levels are illustrated in Fig. 12F. Throughout cervical levels, counts revealed an average of 347 labeled axons in the dCST contralateral to the injection (graph in Figure 12F), with 6 labeled axons in the dCST contralateral to the cortex of origin, and 8 in the dlCST. There were no labeled axons in the ventral column ipsilateral to the injection in the position of the vCST. The number of labeled axons in the dCST remained around 300 through thoracic and lumbar levels, and then fell off at sacral levels.

**Figure 12:**
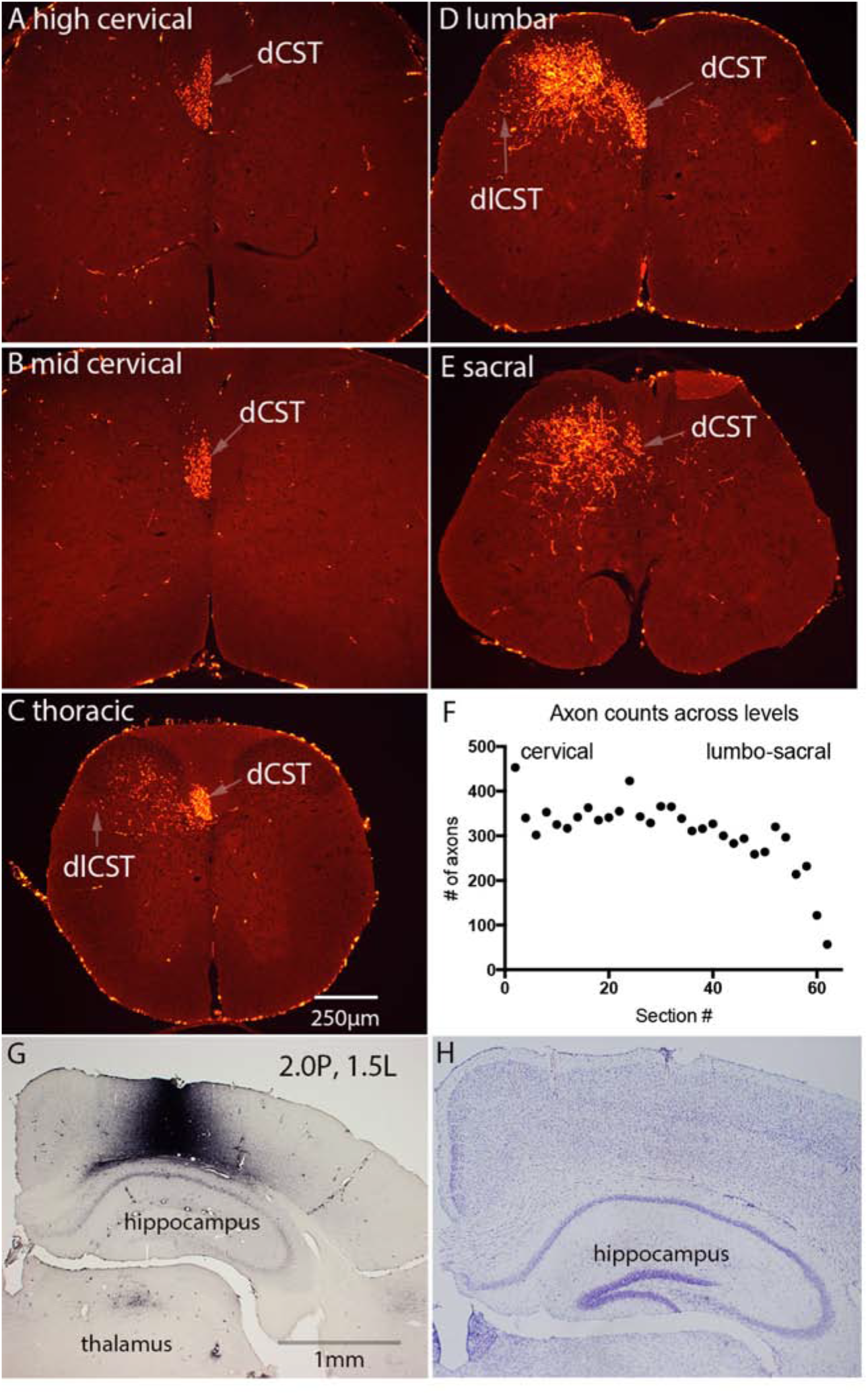
Posterior hindlimb cortex projects selectively to thoracic-sacral levels with terminal arbors located in the deep layers of the dorsal horn. The panels illustrate Cy3-labeled axon arbors at thoracic-sacral levels following a single injection of BDA at 2.0P, 1.5L. A) high cervical level; B) mid cervical; C) low cervical; D) high thoracic; E) mid thoracic; F) Counts of labeled axons in the dCST in sections spaced at 800μm intervals; G) BDA labeling at injection site; H) Cresyl violet stained section at injection site. Abbreviations are as in Fig. 6.

The rostro-caudal distribution of labeled axon arbors was similar to the case illustrated in Figure 11 except that there were almost no labeled arbors in the gray matter at cervical levels. Maximal density of labeled arbors was at lumbar levels, where elaborate axon arbors extended throughout lamina 3-5 of the dorsal horn.

### Projections from the rostral forelimb area

Three mice (0313A-C) received injections targeting the part the dorso-medial frontal cortex that projects to the spinal cord as revealed by retrograde labeling. Target coordinates were 2.8mm anterior and 0.8mm lateral and actual coordinates were 2.6, 2.6, and 3.0mm anterior and 0.7, 0.7, and 0.8mm lateral. Figure 13 illustrates the case (0313C) with the largest number of labeled axons in which the injection site was 2.6mm anterior, 0.7mm lateral (Fig. 13F). Counts revealed an average of 203 labeled axons in the dCST in the first 5 sections of the series (Fig. 13E), 50 labeled axons in the dCST ipsilateral to the cortex of origin, 6 labeled axons in the dlCST and none in the ventral CST. The number decreased progressively moving from rostral to caudal, so that by mid-thoracic levels, only a few labeled axons were present (counts of labeled axons across levels are shown in Figure 13E).

**Figure 13:**
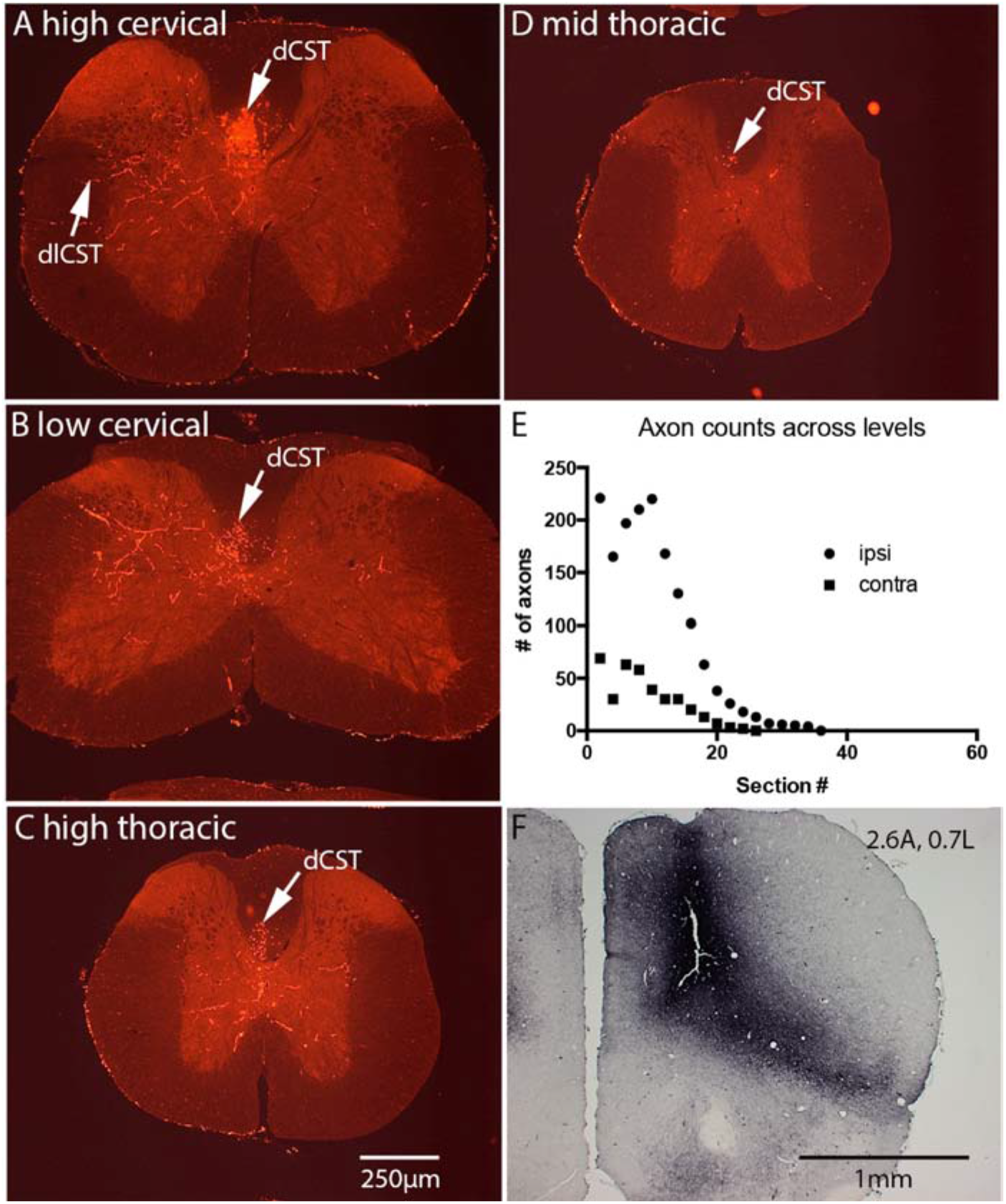
Rostral forelimb area (RFA) projects selectively to cervical-high thoracic levels with terminal arbors located in the intermediate lamina and dorsal part of the ventral horn. The panels illustrate Cy3-labeled axon arbors at cervical-high thoracic levels following a single injection of BDA at 2.6A, 0.7L. A) high cervical level; B) low cervical; C) high thoracic; D) mid thoracic; E) Counts of labeled axons in the dCST in sections spaced at 800μm intervals; F) BDA labeling at injection site.

Labeled axon arbors were present at cervical through mid-thoracic levels and were concentrated in lamina 4-5 of the dorsal horn with a few extending into lamina 7 of the ventral horn. Arborizations were mainly on the side of the labeled tract, but a few axons also extended across the midline.

### Projections from the barrel cortex

Three mice (0408A-C) received injections targeting the dorso-lateral cortex (S2 and the S1 barrel field). Target coordinates were 1.0mm posterior and 3.2mm lateral and 1.1mm deep, and actual coordinates were 1.0, 1.8, and 2.0mm posterior and 3.4, 3.5, and 3.6mm lateral. The distribution of labeled axons in the spinal cord was similar in all 3 cases, and the case with the largest number of labeled axons is illustrated in Figure 14. At high cervical levels, counts revealed 396 and 347 labeled axons in the dlCST in the first 2 sections of the series, with 8 axons in the dlCST and none in the ventral column. The number of labeled axons decreased progressively through the cervical levels and no labeled axons extended beyond low cervical levels.

**Figure 14:**
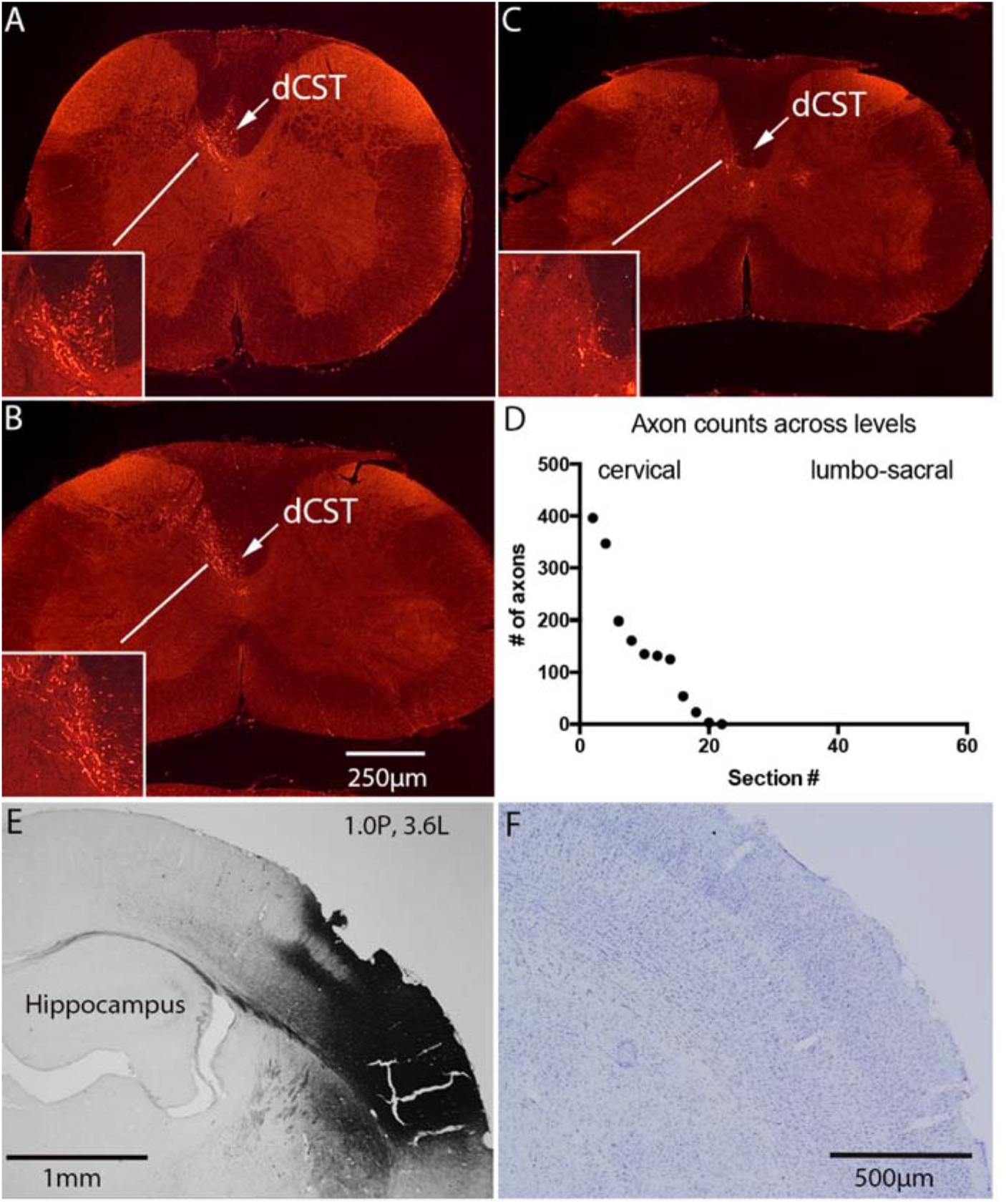
CST neurons in barrel cortex project to high cervical levels and arborizes in the medial intermediate lamina. The panels illustrate Cy3-labeled axon arbors at cervical-high thoracic levels following a single injection of BDA at 1.0P, 3.6L. A) high cervical level; B) mid cervical; C) low cervical; D) Counts of labeled axons in the dCST in sections spaced at 800μm intervals; E) BDA labeling at injection site. F) Cresyl violet stained section at injection site. Abbreviations are as in Fig. 6.

Axon arbors were concentrated in the medial part of lamina 5 of the dorsal horn just lateral and somewhat ventral to the dorsal column. A few labeled axons extended dorsally into the dorsal horn, and a very few extended into the ventral horn. There was a prominent field of labeled axons rostral to the pyramidal decussation in spinal trigeminal *nucleus caudalis* (not shown).

#### Notes on laterality

The majority of CST axons from all locations descended in the main dCST contralateral to the cortex of origin; however, in some mice, there were a significant number of labeled axons in the dCST ipsilateral to the cortex of origin (Figures 9 and 13). This variability did not appear to be systematic, for example, related to location in the cortex, so we attribute this to biological variability. The other aspect of laterality pertains to axons that re-cross at spinal levels to terminate on the side ipsilateral to the cortex of origin. In general, the number of re-crossing axons also varied across cases in a way that did not appear to be systematic, except that re-crossing axons were most common at sacral levels (Figure 11).

## DISCUSSION

Our assessment of the degree to which different parts of the sensorimotor cortex in mice project selectively to different rostro-caudal levels of the spinal cord yielded several conclusions: 1) Projections from the forelimb portion of the sensorimotor cortex are primarily restricted to cervical levels; axons did not extend beyond high thoracic levels; 2) Projections from the hindlimb portion of the sensorimotor cortex pass through cervical regions without extending collaterals into the gray matter; 3) The rostral forelimb area in the dorso-medial frontal cortex projects to cervical through high thoracic levels, and terminates primarily in the intermediate lamina with a few axons extending into the ventral horn; 4) Neurons in the dorso-lateral cortex near the barrel field project exclusively to the high cervical level, arborizing in a small region adjacent to the dorsal column; 5) The degree of bilaterality of terminal arbors varies by spinal level, with the most extensive bilateral projections being at sacral levels; 6) One of the descending pathways seen in other species including rats (the ventral CST) is very sparse in the strains of mice that we studied. 7) Consistent with previous studies in rats (Bareyre et al 2002), CST axons from the medial portion of the sensorimotor cortex terminate primarily in intermediate laminae in the spinal cord with some arbors extending into the ventral horn whereas the lateral portion of the sensorimotor cortex terminates primarily in the dorsal horn.

### Distribution of the cells of origin of the CST in mice

The overall distribution of retrogradely labeled neurons following retro-AAV/Cre or fluorogold injections at C5 is similar to what we have previously described in rats (Nielson et al 2010). The distribution corresponds in general to the motor map of the mouse cortex based on microstimulation (Tennant et al 2011) except that the motor map extends slightly beyond the boundary of the areas containing retrogradely-labeled neurons and the motor map does not include the barrel field in the lateral cortex. Motor maps from microstimulation (Tennant et al 2011) show a rostral forelimb area (RFA) centered at about 2.0 anterior and a larger caudal forelimb area (CFA) the anterior boundary of which is at around 1.0 anterior. Their microstimulation map indicates that the area between RFA and CFA represents the neck and jaw, which are served by lower motoneurons located above C5. CST projections to the motoneurons serving the neck and jaw might not extend to C5, which would account for the lack of retrograde labeling by retro-AAV or fluorogold injections at that level.

The separate group of retrogradely-labeled neurons in the dorso-lateral cortex near the S1 barrel field has been reported in previous studies in rats (Liang et al 2011, Nielson et al 2011). Our results reveal that CST axons from this area terminate selectively at high cervical levels. We are not aware of microstimulation studies targeting this region in rodents. Given the strong projection to *nucleus caudalis* from this region, and the fact that projections to the spinal cord terminate selectively in a small area adjacent to the dorsal horn, this component of CST axons may be involved in modulating sensory processing in the dorsal horn.

### Level-specific projections

BDA injections into the rostral part of the forelimb region lead to extensive labeling of arbors at cervical levels with essentially no extension beyond mid-thoracic levels. With injections into more caudal parts of the forelimb region anterior to bregma, CST axons extended further caudally in the spinal cord. Injections into the hindlimb region posterior to bregma lead to labeling of axons that extend though cervical levels in the dorsal CST largely without extending collaterals. However, injections near the forelimb/hindlimb transition just posterior to bregma led to labeling of arbors at cervical-lumbar and sometimes even sacral levels. Our findings with retro-AAV/Cre indicate that there is some overlap in the distribution of neurons that project to cervical vs. caudal levels, but the absence of double-labeling with injections of retro-AAV/GFP at C5 and retro-AAV/Cre at lumbar levels indicates that there are few if any neurons that project to both levels. This supports conclusions from a previous study that used retro-AAVs with different fluorescent reporters (Wang et al 2018).

### Lamina-specific projections

Previous studies have shown that different parts of the sensorimotor cortex project selectively to different laminae in the spinal cord. This was first shown in hamsters (Kuang & Kalil 1990) and was later confirmed in rats (Bareyre et al 2002). For example, in rats, tracer injections into the “motor cortex” (2mm anterior, 3mm lateral) labeled CST arbors extended throughout the intermediate lamina and into the ventral horn (Bareyre et al 2002). In contrast, following tracer injections into the “sensory cortex” (0.5mm posterior, 4.5mm lateral) labeled CST arbors were restricted to the dorsal horn. The present study confirms the general conclusion for mice; the medial portion of the cortex projects to intermediate lamina and the ventral horn and the lateral portion projects to the dorsal horn. In addition, however, our data reveal through examples that there are actually several different patterns of laminar termination based on point of origin in the cortex.

### CST arbors in the ventral horn; direct innervation of motoneurons?

Previous studies in rats have provided evidence for direct innervation of motoneurons by the CST (Liang et al 1991). Raineteau et al documented extensive projections of CST axons into the cervical ventral horn, and showed close appositions between varicosities on labeled CST axons (putative presynaptic terminals) and retrogradely-labeled motoneuron dendrites (Raineteau et al 2002). Our data reveal that in mice, injections into the medial part of the sensorimotor cortex label elaborate CST arbors in the ventral horn at cervical, lumbar, and sacral levels depending on the injection site and these arbors extend into the portion of the ventral horn containing motoneuron cell bodies and dendrites.

### The ventral CST is sparse in mice

In rats, BDA injections into the sensorimotor cortex label a moderate number of axons in the ventral column ipsilateral to the injection, representing the uncrossed anterior CST that is also present in humans (Brosamle & Schwab 1997). In our previous studies in mice, however, we noted the complete absence of labeled axons in the ventral CST (Steward et al 2004, Steward et al 2008, Zheng et al 2003). These previous studies assessed regenerative growth of CST axons following spinal cord injuries, and for this purpose, mice received a total of 4 BDA injections to blanket most of the sensorimotor cortex rostral and caudal to bregma. In the present study, we again found that there were no labeled axons in the ventral CST in most mice.

Our previous studies in which we did not detect ventral CST axons involved either C57Bl/6 mice or genetically-modified mice with a predominantly C57Bl/6 genetic background. Previous studies of “CST-YFP” mice report YFP-labeled axons in the ventral column that disappear after bilateral pyramidotomy (Bareyre et al 2005). CST-YFP mice are on a mixed genetic background backcrossed to C57Bl/6. Also, one previous study of regenerative growth of CST axons after dorsal hemisection injuries reports increases in the number of CST axons extending along the ventral column following spinal cord injury in mice immunized against Nogo/MAG (Sicotte et al 2003). This study involved SJL/J mice and CST axons were traced by injecting phytohemagglutinin-L (PHA-L) into the sensorimotor cortex. Although it is possible that SJL/J mice have more ventral CST axons than other strains, Sicotte et al did not show examples of labeled vCST axons in un-injured mice. This is important, because we have shown the possibility of artifactual labeling of axons following tracer (BDA) injections at early intervals after spinal cord injury (Steward et al 2007). Indeed, this artifactual labeling occurred after dorsal hemisection injuries as used in Sicotte et al. In sum, although we cannot exclude the possibility that CST axons project via the ventral column in some strains of mice, ventral CST axons are sparse in the strains we have tested (Balb/c, C57/BL, and mice of mixed genetic background). This is relevant for studies of motor recovery after SCI because the ventral CST is one potential source of reinnervation of caudal segments following partial injuries in rats (Weidner et al 2001).

Our results reinforce previous studies (Wang et al 2018) documenting the power of retro-AAV, tissue clearing, and light sheet imaging for obtaining quantitative and complete identification of the components of the CST. Also, our data on the normal projection patterns of CST axons from different parts of the cortex to different levels and laminae of the spinal cord provide novel insights into topographic specificity of normal CST projections, which will aid in identifying differences in projection patterns in mice carrying mutations, in detecting changes in patterns of projection due to regenerative growth following injury (Hollis et al 2016) and form part of the basis for comparisons of CST projection patterns between strains and species.

## Acknowledgements

Thanks to Jamie Mizufuka for superb technical assistance, and to Kiara Quinn for help with editing videos. Supported by NIH grants NS047718 and NS073857 (O.S.), NIH fellowship F31NS070558 (R.W.) and training grants T32GM008620 and EY029596 (R.A.). We gratefully acknowledge generous donations from Cure Medical and Research for Cure as well as other private donations. This study had its roots in a discussion between O. Steward and J. Macklis about the degree to which different parts of the sensorimotor cortex in mice projected selectively to different spinal levels.

